# The origin of the octoploid cloudberry (*Rubus chamaemorus*) genome is the result of multiple and complex polyploidization events

**DOI:** 10.1101/2025.07.07.663483

**Authors:** Marius A. Strand, Thu-Hien To, Ole K. Tørresen, Sarah Fullmer, Morten Skage, Ave Tooming-Klunderud, Simen Rød Sandve, Kjetill S. Jakobsen

## Abstract

We describe a chromosome-level genome assembly from an individual male plant of the cloudberry (*Rubus chamaemorus*). The haplotype-resolved assemblies contain one pseudo-haplotype spanning 1198 megabases and one pseudo-haplotype spanning 1161 megabases. Most of these two assemblies, 93.57% and 96.55% respectively, are each scaffolded into 28 pseudo-chromosomes. Both assemblies show high completeness, with the same BUSCO completeness score of 99.2%. Most BUSCO genes are duplicated in both pseudo-haplotypes, in line with the polyploid nature of the cloudberry genome. The assemblies contain 74,132 and 70,692 predicted protein-coding genes, respectively. Analysis of repetitive sequences classified ∼60% of each haplotype as repeats. Comparative synteny with red raspberry (*Rubus idaeus*) reveals a 4:1 chromosome correspondence, supporting an octoploid origin. Ks distributions and k-mer clustering indicate a fairly recent polyploidization involving one divergent (β) and three closely related (α) subgenomes. Low coverage targeted sequencing data mapped to our assembly link the β-subgenome to a relative of *R. pedatus* or *R. lasiococcus,* while the α-subgenomes might derive from a putative auto-allohexaploid within the main *Rubus* clade. In conclusion, these results indicate that the cloudberry genome arose through multiple hybridization events, including recurrent allopolyploidy and possibly autopolyploidy.

**Significance statement:** Cloudberry cultivation has lagged due to unclear origin and genome structure. We present a haplotype-resolved, chromosome-scale assembly that resolves four homologs per ancestral chromosome into 28 pseudo-chromosomes per haplotype and shows a 4:1 correspondence with red raspberry (*Rubus idaeus*). Genomic analyses indicate a recent polyploidization in which three closely related subgenomes likely derive from a putative hexaploid within the main *Rubus* clade, associated with *R. arcticus* (Arctic raspberry), whereas the fourth, more divergent subgenome is linked to relatives of *R. pedatus* (strawberryleaf raspberry) and *R. lasiococcus* (dwarf bramble). These results clarify cloudberry’s formation and provide a foundation for trait discovery and accelerated breeding.

## Introduction

The cloudberry (*Rubus chamaemorus*) is a circumpolar boreal flowering plant species in the rose family (Urbaniak et al., 2023). Within the rose family (subfamily Rosoideae) *Rubus* is a diverse genus with an estimated number of species ranging from 250–1000 (Huang et al., 2023) found on all continents except Antarctica. Many species of *Rubus* are known for their aggregate fruits such as blackberries, raspberries, dewberries and several hybrids. Aggregate fruits of *R. chamaemorus* are golden-yellow, soft and juicy, rich in vitamin C, ellagic acid, citric acid and anthocyanins, giving it a distinctive tart taste (see Figure 1). They are used in jams, desserts, and liqueurs, and are considered a delicacy, particularly in Nordic countries. Despite great demand, *R. chamaemorus* is not commonly cultivated and is primarily harvested as a wild plant.

**Figure 1:**
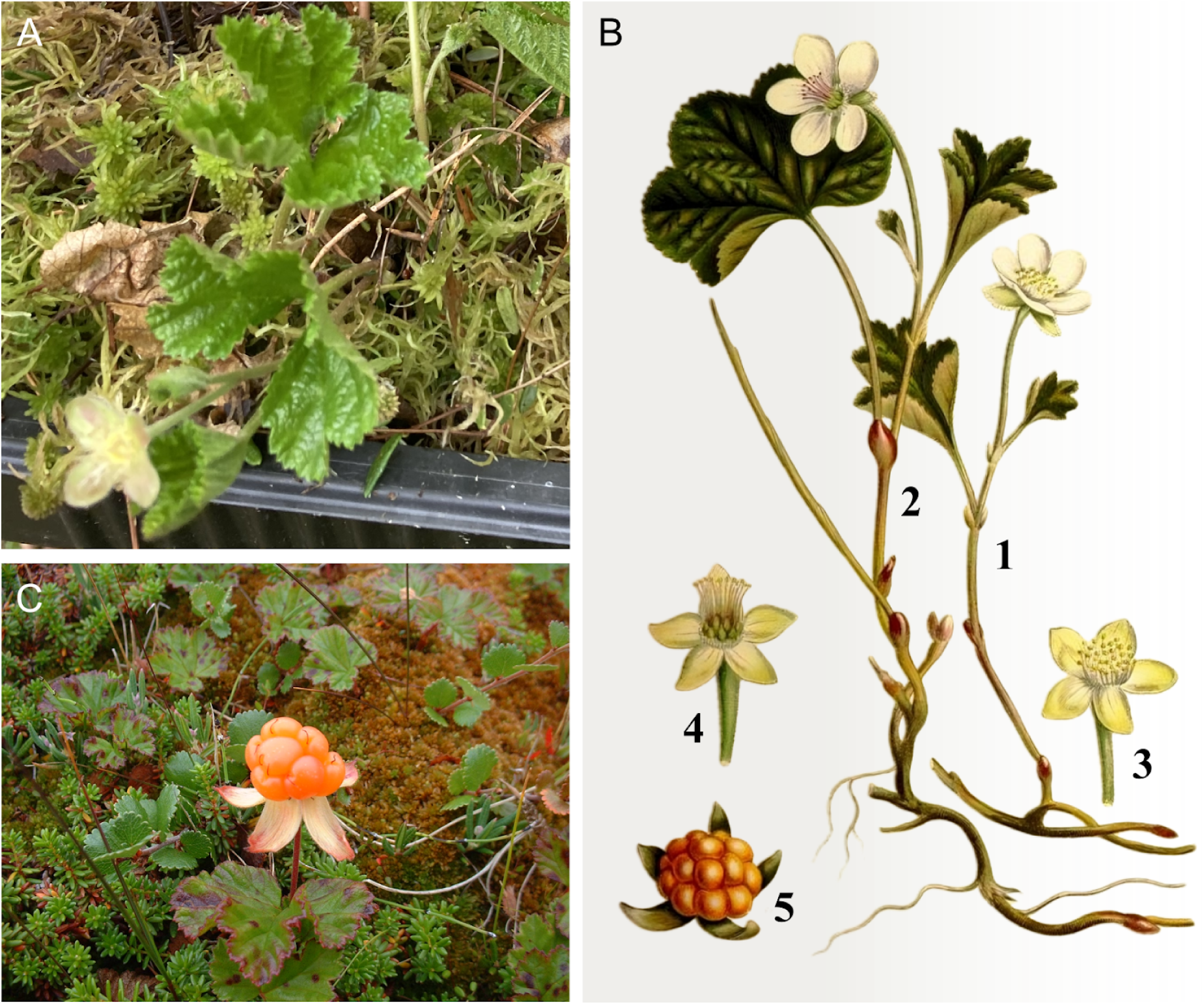
Illustration of dioecy in *Rubus chamaemorus* and the sequenced male specimen. *R. chamaemorus* is dioecious, with male and female flowers borne on separate plants. **A)** Our sample (a male plant) photographed late in the season (photo: Simen R. Sandve). **B)** Historical illustration: (1) male and (2) female plant in bloom; (3) male and (4) female flowers with petals removed; (5) calyx and aggregate fruit. (adapted from *Bilder ur Nordens Flora*, author: C.A.M. Lindman, via https://runeberg.org/nordflor/311.html, public domain). **C)** Female plant with ripe aggregate fruit (image from Wikipedia, author: Philipum, public domain).

Unlike most *Rubus* species, *R. chamaemorus* has biparental (dioecious) reproduction with distinct unisexual individuals, each producing either male or female gametes (see Figure 1B). Dioecy is a strategy to avoid self-fertilization and promote outcrossing. Within *Rubus,* around 60-70% of the species are polyploid ranging from 4x up to 14x (Hummer et al., 2016; Thompson, 1997). *R. chamaemorus* itself is an octoploid with eight genome copies. Evidence from nuclear and chloroplast phylogenies suggests a complex hybrid origin (Carter et al., 2019; Michael, 2006). Michael (2006) found cpDNA support for *R. pedatus* as the maternal ancestor, and nuclear GBSSI-1γ sequences in *R. chamaemorus* revealed two divergent copies: one clustering with *R. lasiococcus*, the other with *R. arcticus*. However, based on ITS data *R. arcticus* was rejected as a possible progenitor. This is consistent with Carter et al. (2019), who placed *R. chamaemorus* in an early-diverging lineage outside the main *Rubus* clade. In addition to interest in its complex polyploidy, *R. chamaemorus* has also received interest for potential cultivar development (Martinussen et al., 2013) as well as regional conservation efforts (Urbaniak et al., 2023).

Despite the potential for cultivar development, no genome assembly has been made available to date. Here, we have used long-read HiFi (PacBio) sequencing combined with long range chromosomal contact mapping by Hi-C, to assemble the first high-quality, haplotype-resolved, chromosome-scale genome of *R. chamaemorus* (Figure 2). The resulting genome consists of two haplotype-resolved assemblies spanning 1198 Mb and 1161 Mb, respectively, with 93.57% and 96.55% of the estimated genome scaffolded into 28 pseudo-chromosomes. Gene annotation identified 74,132 and 70,692 protein-coding genes, respectively. Comparison with other available genome data from *Rubus* species demonstrate that the octoploid *R. chamaemorus* has originated through multiple, and possibly recurrent, hybridizations, leading to the genome architecture of present-day cloudberries. The two haplotype-resolved assemblies provide a genomic foundation for future genetic and evolutionary studies of *R. chamaemorus*.

**Figure 2:**
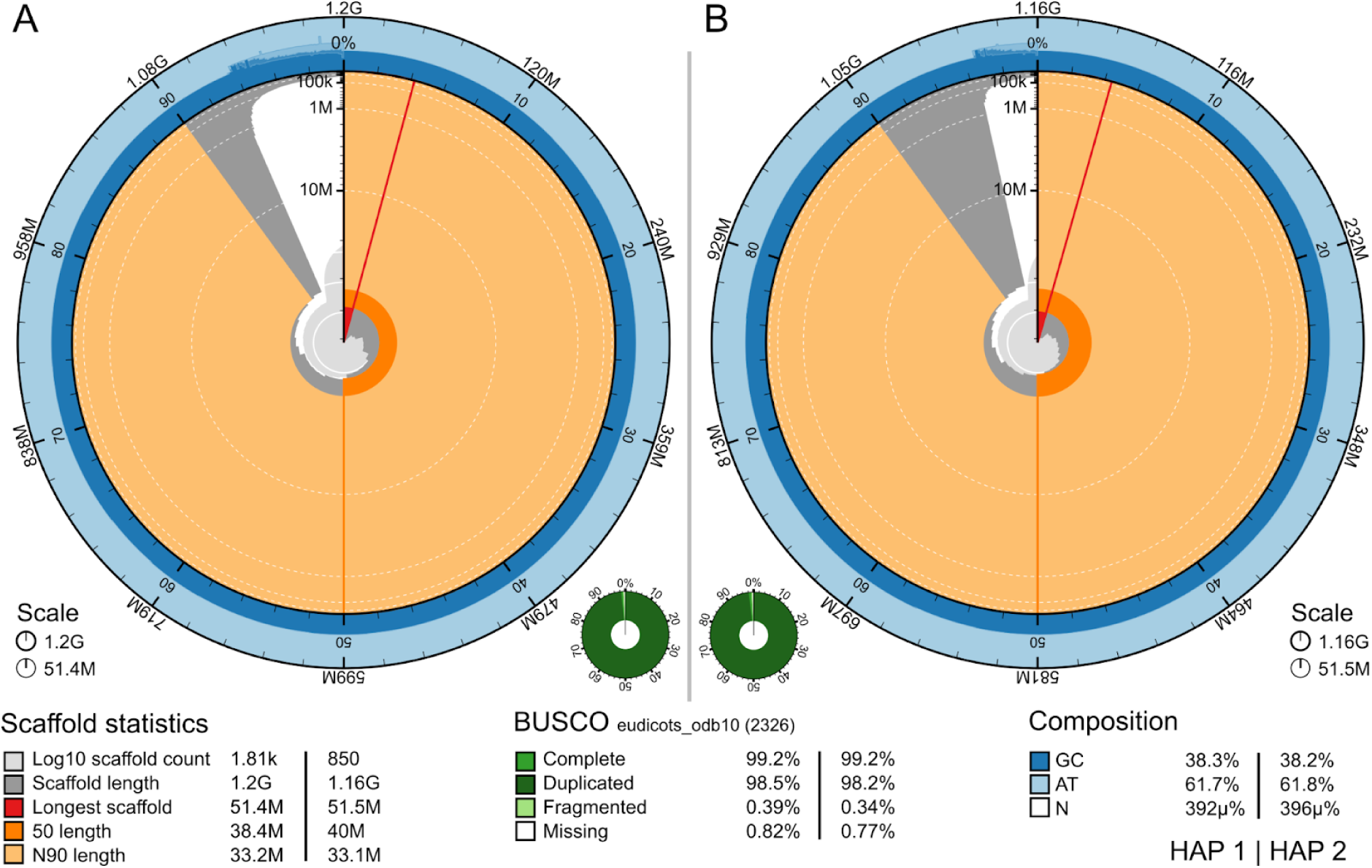
Metrics of the genome assemblies of *R. chamaemorus* hap1 and hap2. **A)** Hap1. **B)** Hap2. The BlobToolKit Snailplots show N50 metrics and BUSCO gene completeness. The two outermost bands of the circle signify GC versus AT composition at 0.1% intervals. Light orange shows the N90 scaffold length, while the deeper orange is N50 scaffold length. The red line shows the size of the largest scaffold. All the scaffolds are arranged in a clockwise manner from the largest to the smallest and are shown in darker gray with white lines at different orders of magnitude, while the light gray shows cumulative count of scaffolds.

## Results

### *De novo* genome assembly and annotation

The genome from *R. chamaemorus* was assembled from a total of 59-fold coverage in Pacific Biosciences single-molecule HiFi long reads and 62-fold coverage in Arima Hi-C reads resulting in two haplotype-separated assemblies. The final assemblies have total lengths of 1198 Mb and 1161 Mb (Table 1 and Figure 2), respectively. Pseudo-haplotypes one (hap1) and two (hap2) have scaffold N50 size of 38.4 Mb and 40.0 Mb, respectively, and contig N50 of 29.4 Mb and 33.1 Mb, respectively (Table 1 and Figure 2). 28 pseudo-chromosomes were identified in both pseudo-haplotypes (ordered by homology to *Rubus idaeus* chromosomes first, and length in hap1 second, and then numbered 1–28, with the homologs in hap2 receiving the same number).

**Table 1:**
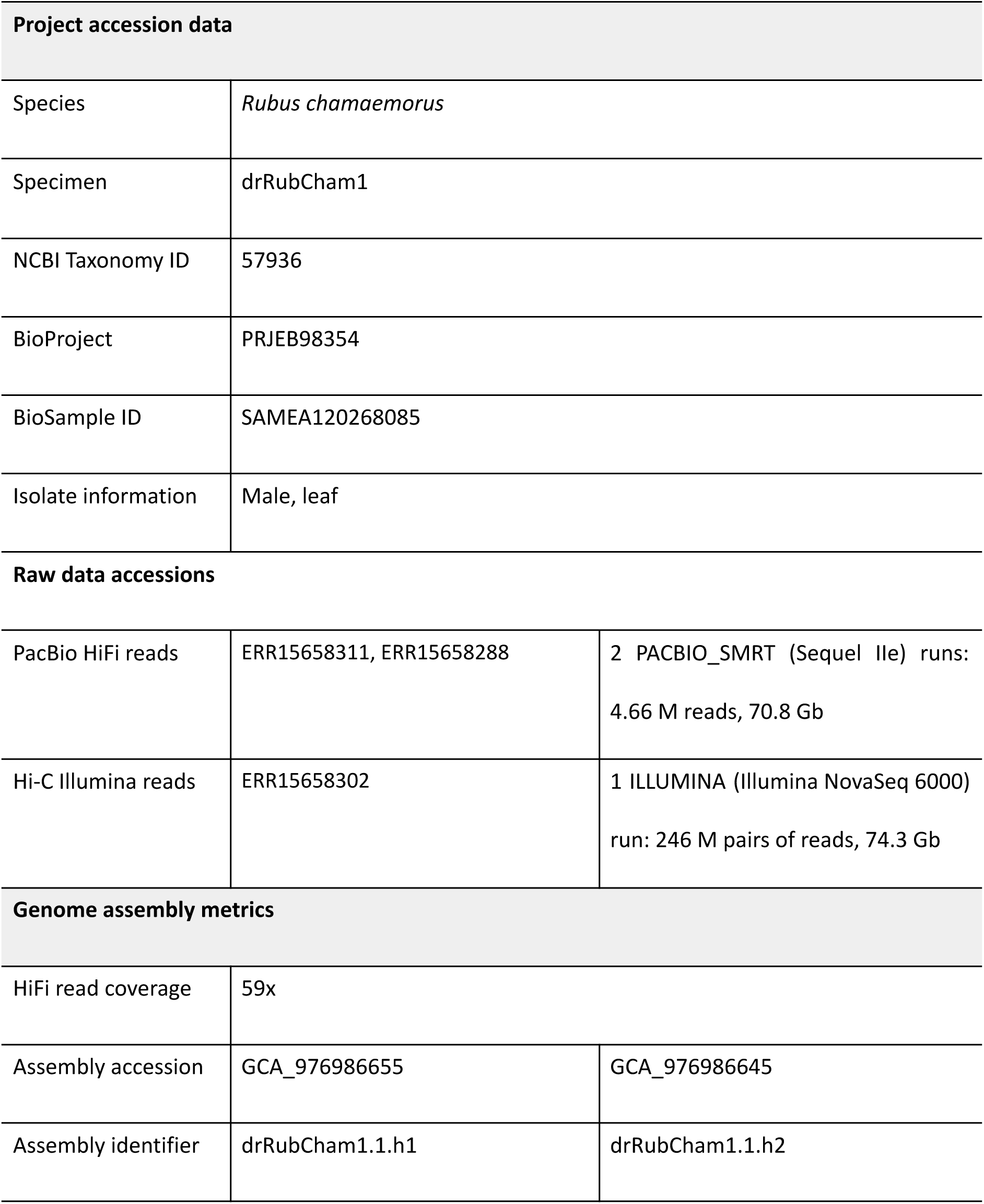
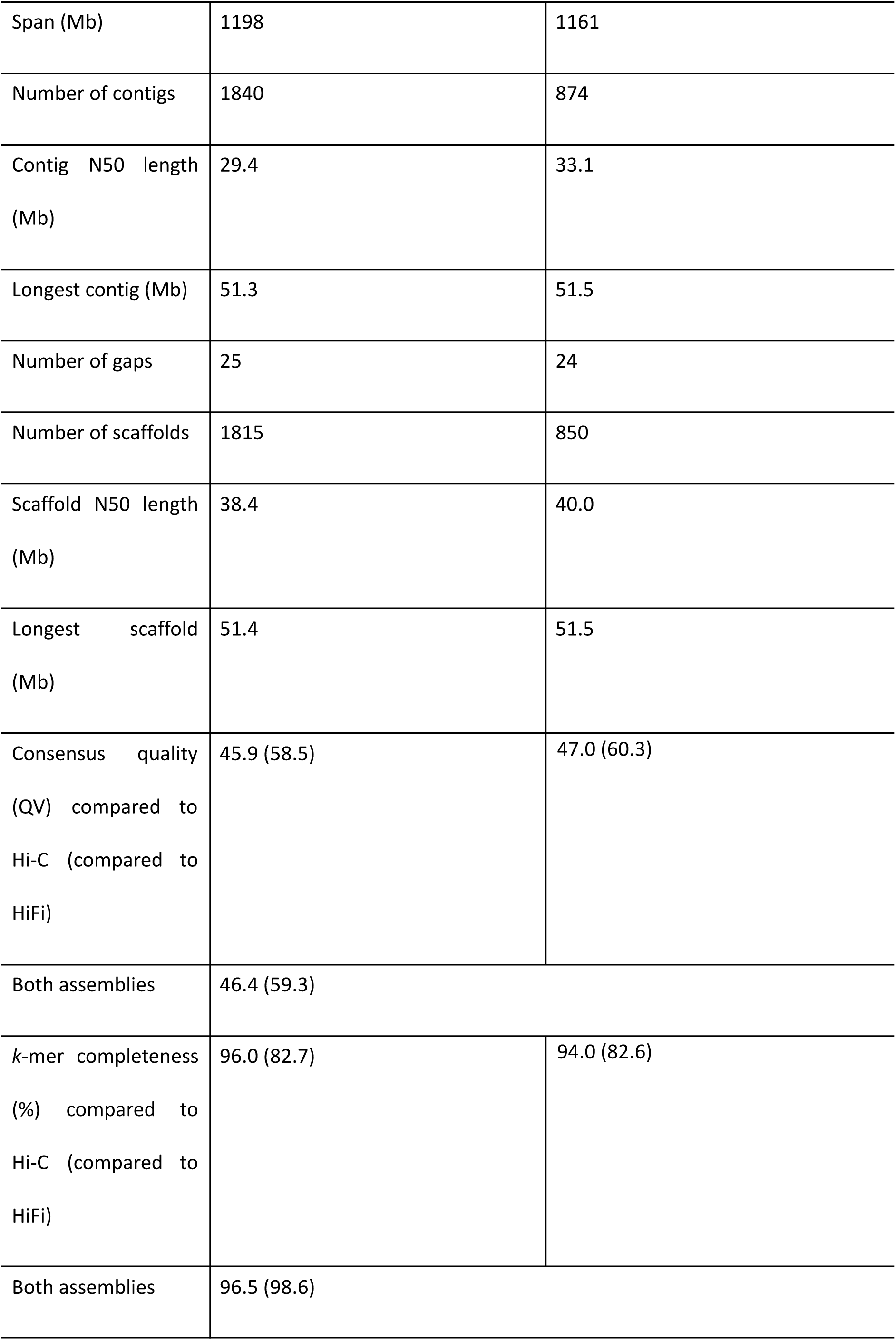
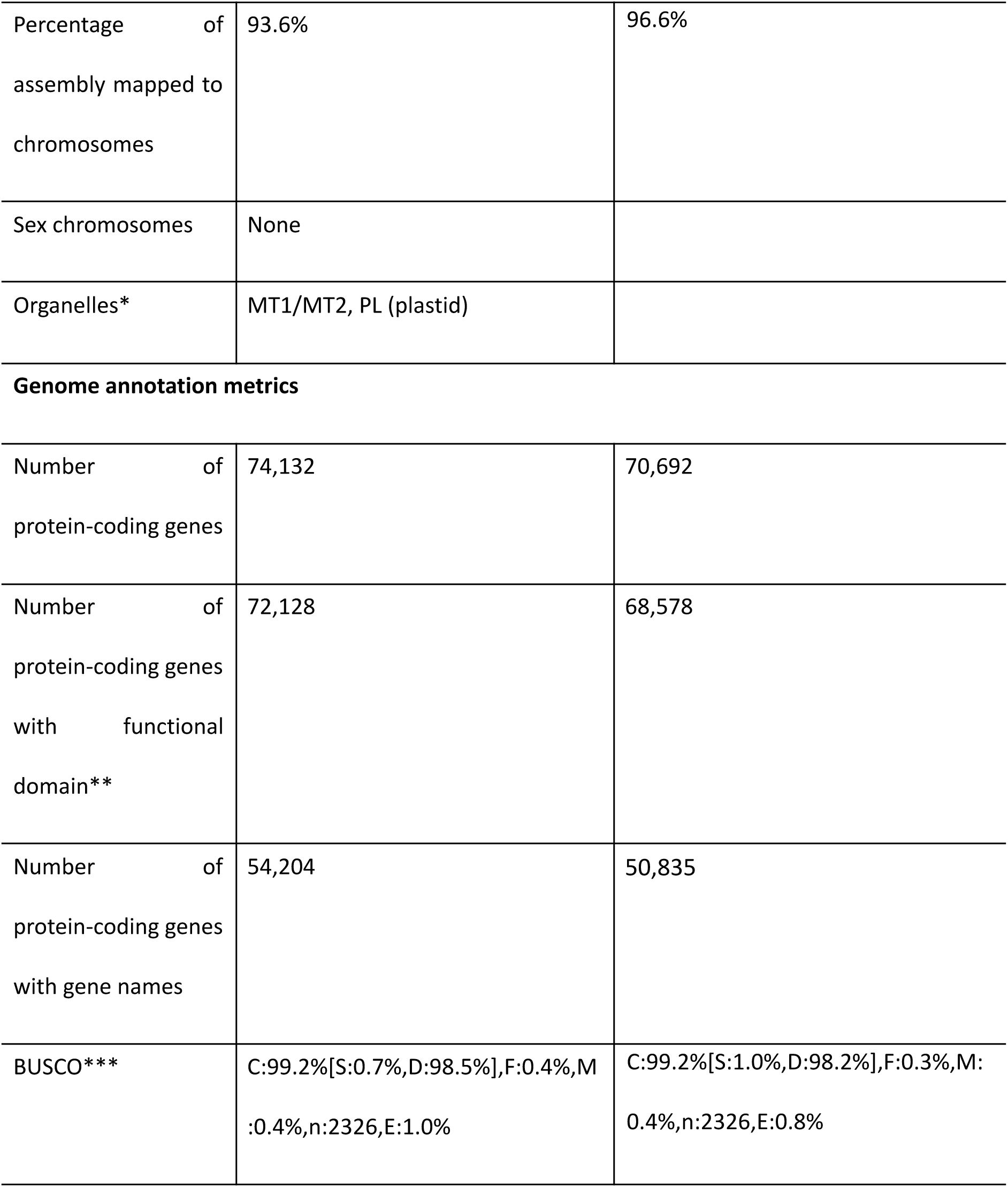
Genome data for *R. chamaemorus*. *Both MT1 and MT2 are partial representations of the full *R. chamaemorus* mitochondria. **Number of genes annotated with a functional domain as found by InterProScan. *** BUSCO scores based on the eudicots BUSCO set using v5.4.7. C = complete [S = single copy, D = duplicated], F = fragmented, M = missing, n = number of orthologues in comparison.

| Project accession data |  |  |
| --- | --- | --- |
| Species | Rubus chamaemorus |  |
| Specimen | drRubCham1 |  |
| NCBI Taxonomy ID | 57936 |  |
| BioProject | PRJEB98354 |  |
| BioSample ID | SAMEA120268085 |  |
| Isolate information | Male, leaf |  |
| Raw data accessions |  |  |
| PacBio HiFi reads | ERR15658311, ERR15658288 | 2 PACBIO_SMRT (Sequel Ile) runs:<br>4.66 M reads, 70.8 Gb |
| Hi-C Illumina reads | ERR15658302 | 1 ILLUMINA (Illumina NovaSeq 6000)<br>run: 246 M pairs of reads, 74.3 Gb |
| Genome assembly metrics |  |  |
| HiFi read coverage | 59x |  |
| Assembly accession | GCA_976986655 | GCA_976986645 |
| Assembly identifier | drRubCham1.1.h1 | drRubCham1.1.h2 |
| Span (Mb) | 1198 | 1161 |
| Number of contigs | 1840 | 874 |
| Contig N50 length (Mb) | 29.4 | 33.1 |
| Longest contig (Mb) | 51.3 | 51.5 |
| Number of gaps | 25 | 24 |
| Number of scaffolds | 1815 | 850 |
| Scaffold N50 length (Mb) | 38.4 | 40.0 |
| Longest scaffold (Mb) | 51.4 | 51.5 |
| Consensus quality (QV) compared to Hi-C (compared to HiFi) | 45.9 (58.5) | 47.0 (60.3) |
| Both assemblies | 46.4 (59.3) |  |
| *k*-mer completeness (%) compared to Hi-C (compared to HiFi) | 96.0 (82.7) | 94.0 (82.6) |
| Both assemblies | 96.5 (98.6) |  |
| Percentage of assembly mapped to chromosomes | 93.6% | 96.6% |
| Sex chromosomes | None |  |
| Organelles* | MT1/MT2, PL (plastid) |  |
| Genome annotation metrics |  |  |
| Number of protein-coding genes | 74,132 | 70,692 |
| Number of protein-coding genes with functional domain** | 72,128 | 68,578 |
| Number of protein-coding genes with gene names | 54,204 | 50,835 |
| BUSCO*** | C:99.2%[S:0.7%,D:98.5%],F:0.4%,M:0.4%,n:2326,E:1.0% | C:99.2%[S:1.0%,D:98.2%],F:0.3%,M:0.4%,n:2326,E:0.8% |

When compared to a k-mer database of the Hi-C reads, hap1 had a k-mer completeness of 96.0%, hap2 of 94.0%, and combined they have a completeness of 96.5%. Further, hap1 had an assembly consensus quality value (QV) of 45.9 and hap2 of 47.0, where a QV of 40 corresponds to one error every 10,000 bp, or 99.99% accuracy compared to a k-mer database of the Hi-C reads (QV 58.5 and 60.3, respectively, compared to a k-mer database of the HiFi reads)(Table 1). When comparing the two pseudo-haplotypes using minimap2, there are 1,775,487 SNP differences (2.098% of the aligned sequence), 149,334 deletions in hap2 compared to hap1 ranging from 1 bp to more than 1000 bp and 149,763 insertions from 1 bp to more than 1000 bp in size (Supplementary Table 2). A total of 74,132 and 70,692 protein-coding genes were annotated in hap1 and hap2, respectively (Table 1).

The Hi-C contact maps show clear separation of chromosomes into homologous sets (Supplementary Figure 1). When sorted by similarity rather than size, each pseudo-haplotype reveals seven distinct groups of four. Repeat masking with RED (Girgis, 2015) identified a high proportion of repetitive content, typical for plant genomes: 60.1% in hap1 and 59.6% in hap2.

Using the phased assembly, we assessed genome-wide synteny between *R. chamaemorus* hap1 and *R. idaeus* (raspberry) – a closely related diploid species with a high-quality chromosome resolved assembly (Martinussen et al., 2013; Price et al., 2023). The synteny analysis revealed a consistent 4:1 chromosome correspondence, supporting an octoploid origin with largely conserved subgenome structure (Figure 3). While most *R. idaeus* chromosomes align directly with four *R. chamaemorus* homologs, several minor deviations indicate historical structural rearrangements. Chromosome 2 in *R. idaeus* shows near-complete synteny with *R. chamaemorus* chromosomes 5–8, but also a small translocation to chromosomes 21–24 (Figure 3B). A similar pattern is seen for *R. idaeus* chromosome 6, which aligns primarily with chromosomes 21–24 but also shows a small translocation to chromosomes 25–28, and a unique translocation from *R. idaeus* chromosome 6 to *R. chamaemorus* chromosome 5. *R. idaeus* chromosome 3 has a smaller central region syntenic across all four homologs of *R. chamaemorus* (chromosomes 9–12). There is also evidence of large inversions. A large central inversion is the simplest explanation for the synteny break between R. idaeus chromosome 2 and chromosomes 7–8, and a similar inversion pattern is observed between chromosome 7 and chromosomes 26–28 (Figure 3B). *R. chamaemorus* chromosome 25 is structurally distinct, lacking this inversion.

**Figure 3:**
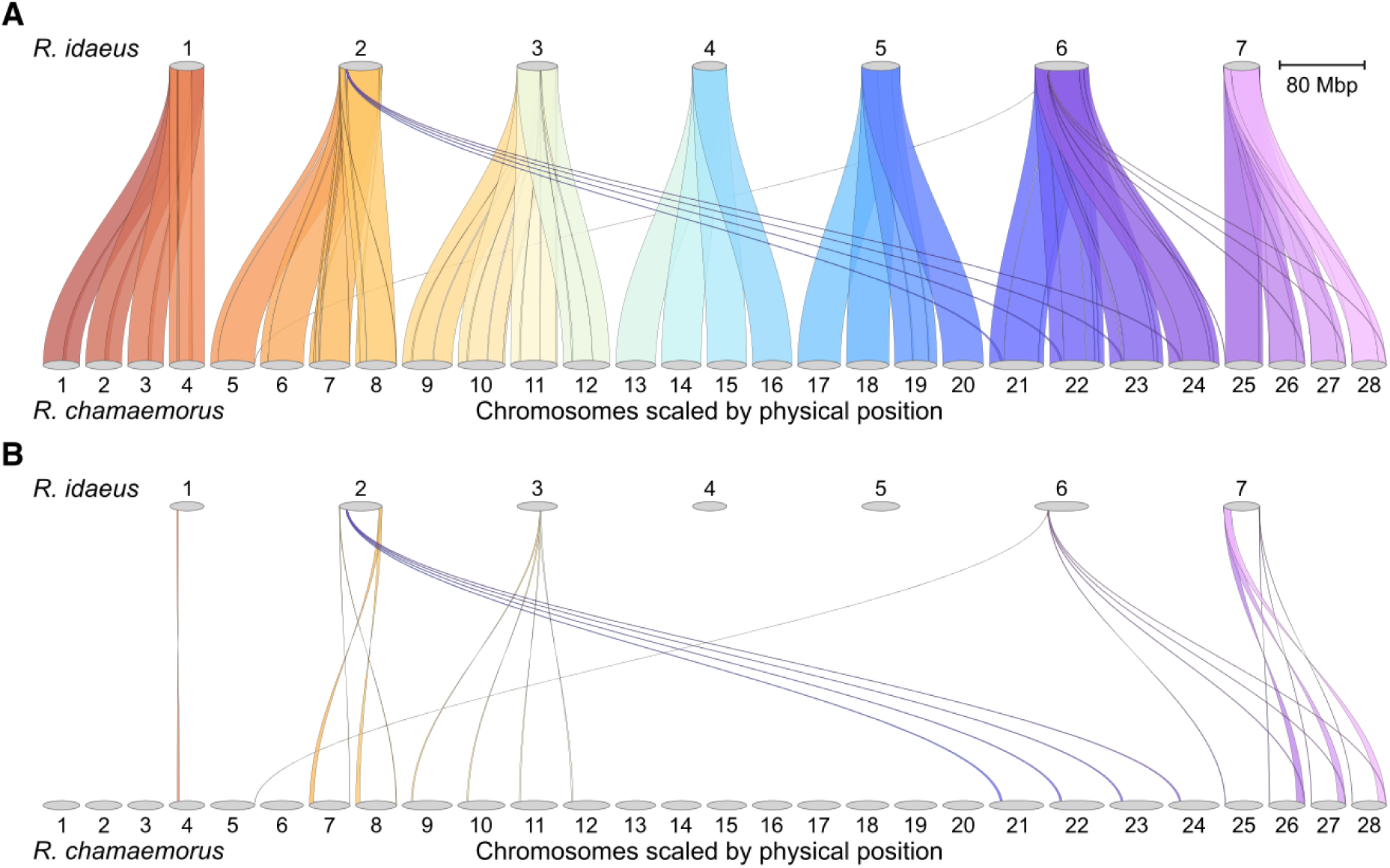
Synteny between *R. chamaemorus* (cloudberry) and *R. idaeus* (raspberry). Both panels are riparian plots generated using GENESPACE. Each *R. idaeus* chromosome aligns with four distinct *R. chamaemorus* hap1 chromosomes, consistent with an octoploid origin. Colors follow a continuous gradient for visual clarity. **A)** Full syntenic alignment. **B)** Alignment plot with conserved collinear blocks manually removed in Inkscape to highlight structural differences (e.g., inversions, translocations).

Hap1 had 99.2% and hap2 99.2% complete BUSCO genes using the eudicots lineage set. Each pseudo-haplotype most commonly has 4 BUSCO copies, consistent with the known octoploid status of *R. chamaemorus* (Table 2). The next most frequent categories are 3 copies, likely reflecting gene loss (fractionation), and >4 copies, which may result from recent small scale duplications or assembly noise. The presence of 2-copy BUSCOs (7.39%) in diploid *R. idaeus* suggests that some genes in the eudicot BUSCO set may not be strictly single-copy within *Rubus*.

**Table 2:** BUSCO duplicates in *R. chamaemorus* compared to *R. idaeus*. Number and percent of BUSCO genes found in *R. chamaemorus* in comparison to *R. idaeus*.

| # BUSCO copies | *R. chamaemorus* hap 1 | *R. chamaemorus* hap 2 | *R. idaeus* |
| --- | --- | --- | --- |
| 0 (Missing) | 10 (0.43) | 10 (0.43) | 32 (1.38) |
| 1 | 26 (1.12) | 32 (1.38) | 2143 (90.76) |
| 2 | 66 (2.84) | 58 (2.49) | 172 (7.39) |
| 3 | 467 (20.08) | 461 (19.82) | 10 (0.43) |
| 4 | 1648 (70.85) | 1659 (71.32) | 1 (0.04) |
| >4 | 109 (4.69) | 106 (4.56) | 0 |
| Total | 2326 | 2326 | 2326 |

To explore the evolutionary history of polyploidization events in *R. chamaemorus*, we first used SubPhaser (Jia et al., 2022) to identify subgenome-specific k-mers and cluster homoeologous chromosomes into putative subgenomes (Figure 4A). The analysis identified one distinct subgenome of 7 chromosomes, hereafter termed the β-subgenome (red, Figure 4A); the remaining 21 chromosomes, lacking such distinction, were collectively grouped as the α-subgenome(s) (grey, Figure 4A).

**Figure 4:**
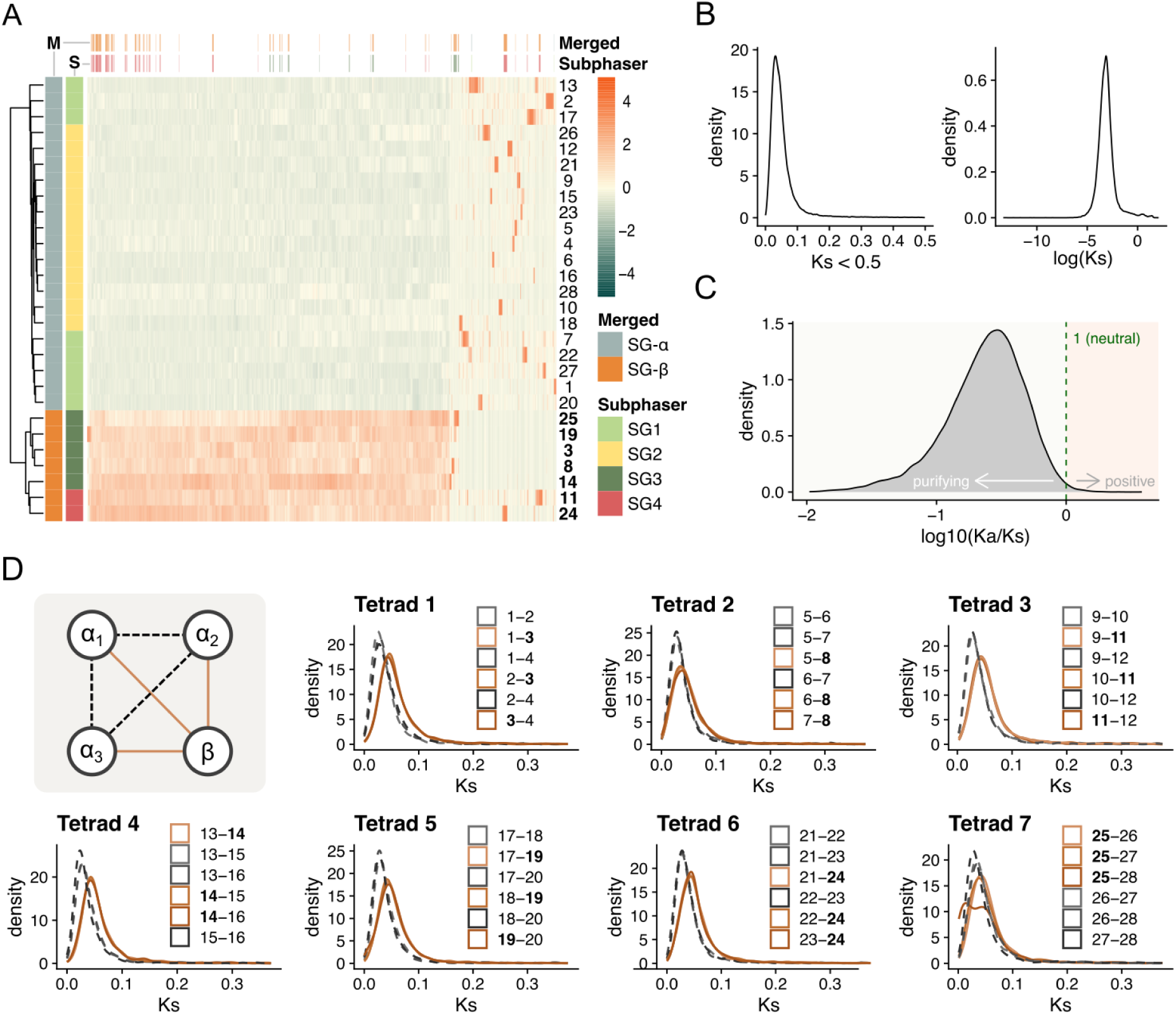
Subgenome structure inferred from k-mer and Ks analyses. **A)** Unsupervised hierarchical clustering of k-mer profiles. Top bar: k-mer subgenome specificity. Side bar: chromosome subgenome assignments. “Merged” labels reflect our inferred α/β subgenomes based on a stricter interpretation of the clustering. Heatmap shows Z-scored relative abundance of 10,000 randomly sampled k-mers. **B)** Ks distribution of segmental duplicates (as defined by doubletrouble) across all chromosomes. Left: linear scale; right: log10-transformed Ks. **C)** Ka/Ks distribution (filtered for Ks > 0.05, Ka/Ks > 0.01; artifact removal), showing most duplicates are under strong purifying selection. **D)** Ks distributions of segmental duplicates within each of the seven syntenic pseudo-chromosome tetrads. The leftmost panel shows pairwise comparisons among the four homologs in each tetrad. Solid orange lines indicate comparisons between each α-homolog and the corresponding β-homolog. Stippled grey lines indicate comparisons among the three α-homologs.

We then estimated Ks distributions using doubletrouble (Almeida-Silva & Van de Peer, 2025). The overall distribution showed a single, narrow peak with only a slight shoulder on the right (Figure 4B). Ka/Ks ratios indicate strong purifying selection across most gene pairs, consistent with functional constraint and subgenome stability (Figure 4C). Stratifying the analysis by tetrad (7 sets of 4 syntenic chromosomes), 6 of the 7 showed a consistent pattern: one chromosome had higher pairwise Ks values relative to the other three (Figure 4D, Supplementary Table 3). These chromosomes correspond to the β-subgenome identified in the k-mer analysis. The 7th tetrad displayed a more complex pattern.

To estimate the time since separation between the α- and the β-subgenomes, as well as within the α-subgenomes, we calculated divergence time (T) using T = Ks / (2 × substitution rate), assuming a constant substitution rate of 4.13 × 10⁻⁹ substitutions/site/year. The overall trend (Supplementary Figure 4) shows that the α- and β-subgenomes diverged approximately 5.83 Ma (95% CI: 5.80–5.87; median Ks = 0.0481), while the median duplication age within the α-subgenomes is around 4.17 Ma (95% CI: 4.14–4.21; median Ks = 0.0345); roughly 1.7 million years later. Across tetrads, Ks values between α-subgenome chromosomes vary moderately but show no consistent clustering pattern, suggesting that divergence within the α-subgenome is relatively symmetric and lacks a clear hierarchy. Tetrad 7 is an exception to the other tetrads, showing greater variability and less internal consistency especially between chromosomes 25, 26 and 28.

Given at least one hybridization event in the origin of *R. chamaemorus*, we assessed whether reads from candidate *Rubus* taxa map preferentially to α or β*. R. pedatus* and *R. lasiococcus* have previously been proposed as progenitor species of *R. chamaemorus* (Carter et al., 2019; Michael, 2006), while *R. arcticus* shares a GBSSI-1γ copy with *R. chamaemorus (Michael, 2006)*. Although genome assemblies are not available for these species, limited sequencing data is available (through Carter et al. (2019) and Kates et al. (2024); see Material and Methods). To minimize mapping ambiguity among closely related α subgenomes, we retained only the longest α representative per tetrad and quantified high-confidence alignments across α and β. Using each phylogenetic tree as a scaffold, we mapped log₂ coverage bias to the reduced genome (full species lists and complete trees in Supplementary Figures 5–8).

In the Carter dataset, we observe the strongest β bias within the *R. pedatus–R. lasiococcus* clade, with weak β bias or near neutrality in other early-diverging *Rubus* lineages (Figure 5A; group 2). The main *Rubus* clade is predominantly α-biased in Carter, with a few neutral species nested within otherwise α-biased subclades (Supplementary Figures 5–6). When the signal is examined by tetrad, tetrad 7 (chr25–28; chr25 vs. 26) behaves idiosyncratically: *R. idaeus* exhibits an anomalously strong β bias (Figure 5B), and several otherwise weakly biased species show the same pattern specifically for this tetrad. Neither *R. idaeus* nor these other species show comparable β bias in any other tetrads.

**Figure 5:**
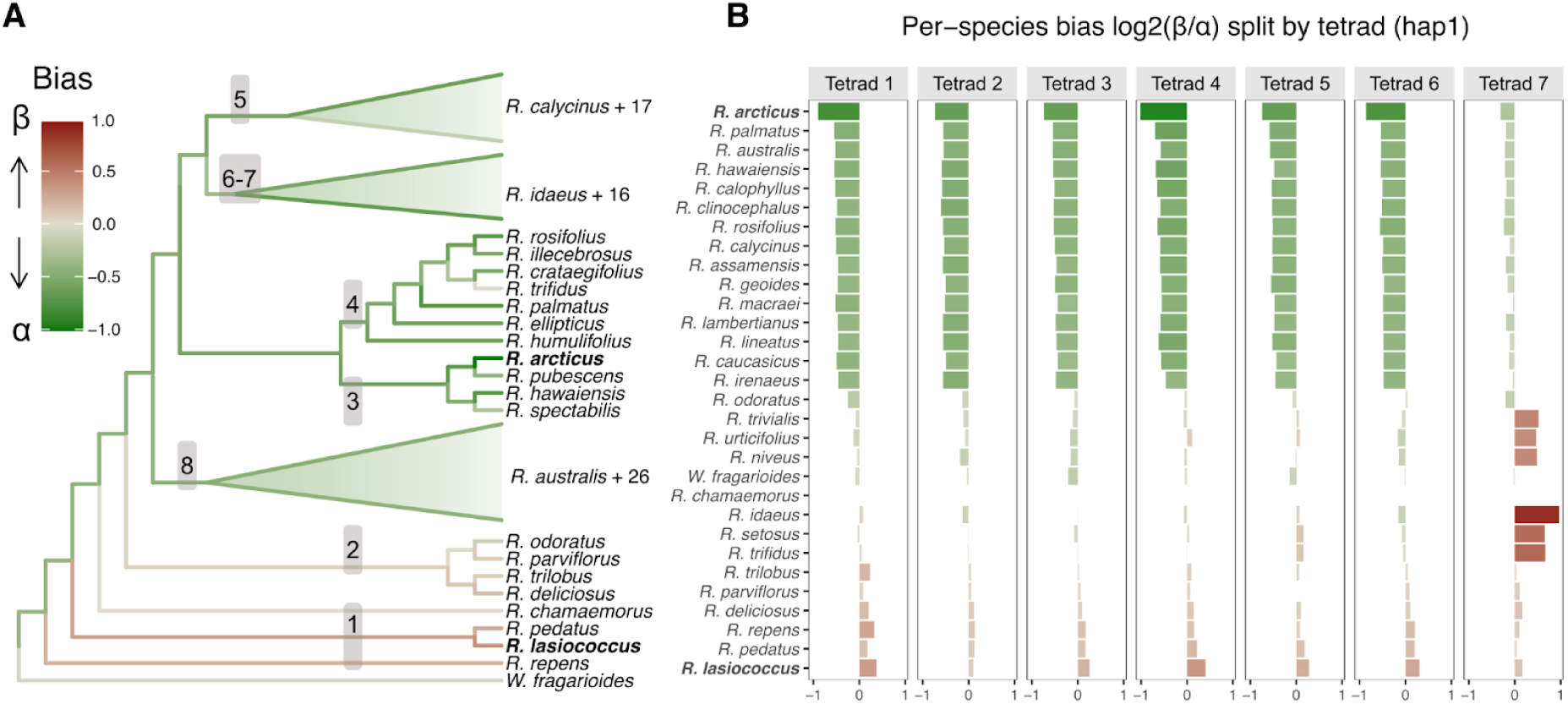
High-confidence read mappings (MAPQ *≥* 30) from Carter et al. (2019) depicting bias of *Rubus* species mapped to a reduced *α* and *β* subgenome set of *R. chamaemorus*. Reads from each species are mapped to *R. chamaemorus* haplotype one (hap1). Log₂ of the ratio of high-confidence reads mapped to subgenome β over subgenome α, with bias shifted so that *R. chamaemorus* is at 0. A value of 1 indicates a two-fold mapping bias toward subgenome β. Chromosomes from subgenome α: 1, 5, 9, 13, 17, 21, 26; from subgenome β: 3, 8, 11, 14, 19, 24, 25. **A)** *R. chamaemorus* centered bias (median across tetrads) projected onto the ASTRAL-II exon all-taxa tree; the tree is collapsed to show only relevant lineages. (Numbers denote Carter et al. groups; Fig. 2). The most α-biased and β-biased species are marked in bold. **B)** Bias stratified by chromosome pair (tetrad). Species are ordered from most α-biased (most negative) to most β-biased (most positive). Only the 15 most α-biased and the 15 least α-biased species are shown.

In the Kates dataset, overall β bias is stronger than in Carter, shifting groups 5–8 to be neutral/slightly β-biased, with *R. pedatus* showing the strongest β bias and *R. arcticus* and associated clades (3–4) retaining their α-biased status (Supplementary Figures 7–8). While the overall bias in Kates is either neutral or β-biased, tetrad 6 stands out with its strong and consistent α-bias. Unlike in Carter there are spikes of β-bias in tetrads 4 and 5, as well as tetrad 7, which might be a result of fewer and less informative loci for *Rubus* in this dataset.

### Subgenome evolution scenarios

Together, Figures 3–5 constrain the set of plausible polyploidization pathways leading to the octoploid *R. chamaemorus*. The subgenomes form two main lineages, α and β, which started diverging ∼5.83 Ma. Within α, there are three sets of 7 (α_1_–α_3_), but k-mer composition and Ks profiles do not resolve them as distinct subgenomes. We therefore treat α_1_–α_3_ as interchangeable topologically and use the numbering only to track hypothetical compositions. Under the criterion that only one subgenome is distinguishable, the 15 theoretical topologies collapse to four end-state topologies (Figure 6A; putative scenarios depicted in Figure 6 B–F).

**Figure 6.**
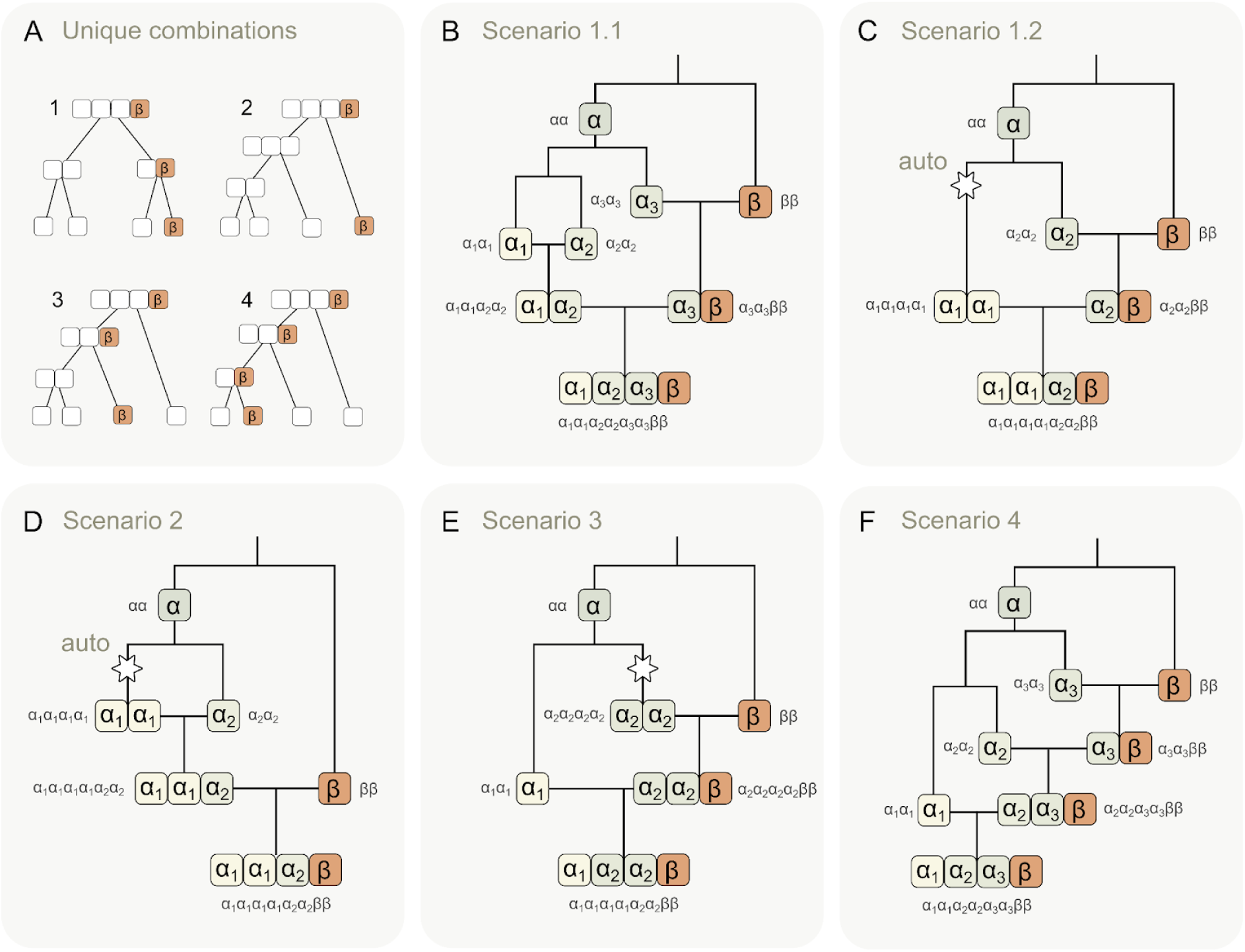
Scenarios for subgenome origin and polyploidization. **A)** When only one of four subgenomes is distinguishable, the 15 possible octoploid topologies collapse into four equivalence classes; One where the intermediates are tetraploids (Scenario 1), and 3 where a hexaploid is an intermediate (Scenarios 2–4). **B)** Scenario 1.1: three successive hybridizations (α_3_ × β and α_1_ × α_2_, then α_3_β × α_1_α_2_) yield four distinct subgenomes. **C)** Scenario 1.2: one autotetraploidization of α_1_ (α_1_ → α_1_α_1_), followed by α_2_ × β and α_1_α_1_ × α_2_β, yields three distinct subgenomes. **D–F)** Scenarios 2–4: routes via a hexaploid intermediate that differ only in the order of β incorporation. Scenario 2: β joins after formation of α_1_α_1_α_2_. Scenario 3: β first merges with tetraploid α_2_α_2_ to form a hexaploid, which then merges with α_1_. Scenario 4: β sequentially merges with three α lineages, forming a tetraploid, then a hexaploid, then the octoploid.

Figures 6B and 6C both show topology 1, in which an octoploid forms via two tetraploids, differing only by the mechanism of α polyploidization. In Scenario 1.1 (Figure 6B), three successive hybridizations; α_3_ × β and α_1_ × α_2_, followed by α_3_β × α_1_α_2_; yield four distinct subgenomes. In Scenario 1.2 (Figure 6C), α_1_ first undergoes autotetraploidization (α_1_ → α_1_α_1_), α_2_ hybridizes with β, and the merger α_1_α_1_ × α_2_β produces an octoploid with three distinct subgenomes.

Figures 6D–F depict topologies 2–4, each proceeding through a hexaploid intermediate and differing only in the order of β incorporation. Scenario 2 (Figure 6D) begins with autotetraploidization of α_1_, followed by hybridization with α_2_ (α_1_α_1_ × α_2_), and a subsequent merger with β (α_1_α_1_α_2_ × β), yielding an octoploid with three distinct subgenomes. In Scenario 3 (Figure 6E), β first merges with a tetraploid α_2_α_2_ to form a hexaploid, which then merges with α_1_. In Scenario 4 (Figure 6F), β merges with one α (α_3_) to form a tetraploid, then with a second α (α_2_) to form a hexaploid, and finally with a third α (α_1_) to produce an octoploid.

## Discussion

Polyploidy is pervasive in *Rubus*, with 60–70% of species possessing chromosome numbers 4n or higher (Chen et al., 2020; Hummer et al., 2016; Thompson, 1997). Comparative synteny between *R. chamaemorus* and *R. idaeus* reveals a highly collinear subgenome structure, with four homologs corresponding to each of the seven basal chromosomes typical of *Rubus*.

K-mer clustering and Ks analyses suggest an evolutionary history in which three of the four subgenomes originate from closely related species or populations (α), while the fourth is more divergent (β) (Figure 4). The lack of resolvable α subgenome structure suggests limited divergence among α copies, with subsequent homoeologous exchange erasing most differences, including accumulated repeats and transposable elements. Although such erasure permits multiple histories, Scenario 2, outlined in figure 6D, is the most parsimonious: an α autotetraploid merges with a closely related α diploid to form an auto-allohexaploid with three α copies, which then merges with the more divergent β diploid.

Using a reduced reference comprising one α-subgenome set (longest chromosome) and the β-subgenome, we mapped short reads from Carter et al. (2019) and Kates et al. (2024) to estimate α–β mapping bias. Since Carter et al. specifically targets *Rubus* and uses ∼1,000 low-copy nuclear genes, in contrast to Kates et al. who use 100 loci across the nitrogen-fixing clade only incidentally covering *Rubus*, we defer to Carter for the most accurate phylogeny of *Rubus* and the most reliable results, treating Kates as a cross-check for mapping bias. Both datasets recover the same extremes: strongest β affinity in the *R. pedatus*–*R. lasiococcus* lineage and strongest α affinity within the main *Rubus* clade, specially Carter et al. (2019) groups 3–4 with *R. arcticus*, where α-bias persists in Kates despite the overall β shift.

Although it has been proposed that both R. pedatus and *R. lasiococcus* are progenitors of *R. chamaemorus* (Michael, 2006), our results indicate they may only account for the β subgenome (Figure 5A). Based on the available chloroplast trees, it is likely the maternal ancestor is more closely related to *R. pedatus* (Carter et al., 2019; Michael, 2006). *R. arcticus* (group 3) exhibits the strongest α bias in Carter, followed by *R. palmatus in* the adjacent group 4 (Figure 5). Taken together, these patterns are consistent with an unsampled or extinct α-donor lineage along the stem leading to groups 3–4, rather than from any single species present in the datasets.

Our estimate of ∼5.83 Ma as the start of α–β divergence, which preceded the polyploidization events leading to *R. chamaemorus,* should be regarded as highly uncertain, as the substitution rates for herbaceous perennials found by Kay et al. (2006) ranges from 1.72 × 10^−9^ to 8.34 × 10^−9^ substitutions/site/year. Using the low estimate, the split between α- and β-subgenomes comes out to 14.1 Ma, which is more consistent with the time estimates of the split between *R. lasiococcus*–*R. pedatus* and the main *Rubus* clade estimated by Carter et al. (2019). It is also possible, given an evolutionary setting with frequent hybridizations and homoeologous exchanges over time, that divergence is underestimated.

Tetrad 7 (chromosomes 25–28) consistently departs from the otherwise clear α–β pattern across analyses. Only the K-mer analysis cleanly assigns chromosome 25 to β and chromosomes 26–28 to α, indicating unique sequence content on chromosome 25. In contrast, pairwise Ks within the tetrad do not follow the typical α–β divergence pattern: comparisons involving chromosome 25, especially with chromosome 28, are bimodal (Supplementary Figure 3), suggesting a mosaic history. Mapping-bias analysis further highlights this anomaly, with several species, most notably *R. idaeus*, showing a stark shift toward β only in this tetrad (Figure 5B). Hi-C and collinearity analyses provide no clear evidence of misjoins (Figure 3 and Supplementary Figure 1), and chromosome 25 uniquely lacks the large inversion present on chromosomes 26–28 (Figure 3B), which suggests a biological rather than assembly cause. Although data artifacts cannot be ruled out, these consistent irregularities across methods point to a complex evolutionary history for Tetrad 7.

Resolving these questions will require high-quality genome assemblies of the putative relatives of *R. chamaemorus*; including *R. pedatus*, *R. lasiococcus*, *R. arcticus*, and related taxa. More broadly, such assemblies are essential for disentangling polyploid origins, identifying subgenome contributors, and enabling accurate comparative genomics across complex plant lineages.

The presented *R. chamaemorus* assemblies are a resource that can support cultivar development. Knowledge of the polyploid genome structure, combined with a high-quality assembly and annotation, can accelerate breeding in *R. chamaemorus*, where conventional approaches are limited by dioecy and slow maturation (Martinussen et al., 2013). One promising target trait is primocane-fruiting (fruiting on first-year stems), which occurs naturally in many *Rubus* species, including raspberries and some blackberries (Jennings, 2018). In blackberry, the diploid ‘Hillquist’ (*R. argutus*) enabled identification of candidate flowering-time genes for this trait (Brůna, Aryal, et al., 2023). Similar strategies may be explored in *R. chamaemorus*, using genomic data to identify flowering-related loci and reduce the long breeding cycle.

## Material and Methods

### Sample acquisition and DNA extraction

The sample was derived from an adult male plant collected in a forested area of Ås, Norway (59.670983, 10.789649) on the 29th of May 2022 (Figure 1A). Sampling of this plant material does not require a specific permit under Section 15 of the Norwegian Nature Diversity Act, provided that it does not threaten the viability of the population and is not otherwise restricted.

DNA isolation for PacBio long read sequencing was performed using Circulomics Nanobind CBB BIG DNA kit and protocol according to manufacturer’s recommendations, including treatment with EtOH removal buffer (Circulomics, now PacBio company). Quality check of amount, purity and integrity of isolated DNA was performed using Qubit BR DNA quantification assay kit (Thermo Fisher), Nanodrop (Thermo Fisher), and Fragment Analyser (DNA HS 50kb large fragment kit, Agilent Tech.).

### Library preparation and sequencing for *de novo* assembly

Before PacBio HiFi library preparation, DNA was purified an additional time using AMPure PB beads (1:1 ratio). Purified HMW DNA was sheared into an average fragment size of 15-20 kbp large fragments using the Megaruptor3 (Diagenode). A HiFi library was prepared following the PacBio protocol for HiFi library preparation using the SMRTbell® Prep Kit 3.0. The final HiFi library was size-selected with a 10 kbp cut-off using a BluePippin (Sage Sciences) and sequencing was performed by the Norwegian Sequencing Centre on a PacBio Sequel IIe instrument (Pacific Biosciences Inc). The library was sequenced on two 8M SMRT cells using a Sequel II Binding kit 3.2 and Sequencing chemistry v2.0.

A Hi-C library was prepared using the Arima High Coverage HiC kit (Arima Genomics), following the manufacturer’s recommendations and starting with 1050 mg leaf tissue. Chromatin was crosslinked prior to nuclei isolation as written in the user guide for plant tissues (Doc A160163 v01). Final library quality was quantified using a Kapa Library quantification kit for Illumina (Roche Inc.). The library was sequenced with other libraries on an Illumina NovaSeq 6000 (Illumina Inc) with 2 × 150 bp paired end mode at the Norwegian Sequencing Centre (https://www.sequencing.uio.no).

### Genome assembly and curation, annotation and evaluation

A full list of relevant software tools and versions is presented in supplementary table 1. We assembled the species using a pre-release of the EBP-Nor genome assembly pipeline (https://github.com/ebp-nor/GenomeAssembly). KMC (Kokot et al., 2017) was used to count k-mers of size 32 in the PacBio HiFi reads, excluding k-mers occurring more than 10,000 times. GenomeScope (Ranallo-Benavidez et al., 2020) was run as part of the pipeline on the k-mer histogram output from KMC and was included in the methods for completeness, but as it does not support ploidy above 6, we do not report its results. Ploidy level was investigated using Smudgeplot (Ranallo-Benavidez et al., 2020). HiFiAdapterFilt (Sim et al., 2022) was applied on the HiFi reads to remove possible remnant PacBio adapter sequences. The filtered HiFi reads were assembled using Hifiasm (Cheng et al., 2021) with Hi-C integration resulting in a pair of haplotype-resolved assemblies, pseudo-haplotype one (hap1) and pseudo-haplotype two (hap2). Unique k-mers in each assembly/pseudo-haplotype were identified using meryl (Rhie et al., 2020) and used to create two sets of Hi-C reads, one without any k-mers occurring uniquely in hap1 and the other without k-mers occurring uniquely in hap2.

K-mer filtered Hi-C reads were aligned to each scaffolded assembly using BWA-MEM (Li, 2013) with `-5SPM` options. The alignments were sorted based on name using samtools (Li et al., 2009) before applying samtools fixmate to remove unmapped reads and secondary alignments and to add mate score, and samtools markdup to remove duplicates. The resulting BAM files were used to scaffold the two assemblies using YaHS (Zhou et al., 2023) with default options. FCS-GX (Astashyn et al., 2024) was used to search for putative contamination and contaminated sequences were removed. The mitochondrion was searched for in contigs and reads using Oatk (Zhou et al., 2024). The assemblies were manually curated using PretextView and Rapid curation 2.0. Chromosomes were identified by mapping to *Rubus idaeus* JASCWY01 (Price et al., 2023) and *Fragaria vesca* v4 (Edger et al., 2018) in addition to inspecting the Hi-C contact map in PretextView. We annotated the genome assemblies using a pre-release version of the EBP-Nor genome annotation pipeline (https://github.com/ebp-nor/GenomeAnnotation). First, AGAT (https://zenodo.org/record/7255559) `agat_sp_keep_longest_isoform.pl` and `agat_sp_extract_sequences.pl` were used on the *Arabidopsis thaliana* (TAIR10.1, GCF_000001735.4) genome assembly and annotation to generate one protein (the longest isoform) per gene. Miniprot (Li, 2023) was used to align the proteins to the curated assemblies. UniProtKB/Swiss-Prot (UniProt Consortium, 2023) release 2023_02 in addition to the Viridiplantae part of OrthoDB v11 (Kuznetsov et al., 2023) were also aligned separately to the assemblies. RED (Girgis, 2015) was run via redmask (https://github.com/nextgenusfs/redmask) on the assemblies to mask repetitive areas. GALBA (Brůna, Li, et al., 2023; Buchfink et al., 2015; Hoff & Stanke, 2019; Li, 2023; Stanke et al., 2006) was run with the *Arabidopsis* proteins using the miniprot mode on the masked assemblies.

The `funannotate-runEVM.py` script from Funannotate was used to run EvidenceModeler (Haas et al., 2008) on the alignments of TAIR10.1 proteins, UniProtKB/Swiss-Prot proteins, Viridiplantae proteins and the predicted genes from GALBA. The resulting predicted proteins were compared to the protein repeats that Funannotate distributes using DIAMOND blastp and the predicted genes were filtered based on this comparison using AGAT. The filtered proteins were compared to the UniProtKB/Swiss-Prot release 2023_02 using DIAMOND (Buchfink et al., 2015) `blastp` to find gene names and InterProScan (Jones et al., 2014) was used to discover functional domains. AGATs `agat_sp_manage_functional_annotation.pl` was used to attach the gene names and functional annotations to the predicted genes. EMBLmyGFF3 (Norling et al., 2018) was used to combine the fasta files and GFF3 files into an EMBL format for submission to ENA.

All the evaluation tools have also been implemented in a pipeline, similar to assembly and annotation (https://github.com/ebp-nor/GenomeEvaluation). Merqury (Rhie et al., 2020) was used to assess the completeness and quality of the genome assemblies by comparing them to the k-mer content of both the Hi-C reads and PacBio HiFi reads. BUSCO (Manni et al., 2021) was used to assess the completeness of the genome assemblies by comparing against the expected gene content in the eudicots lineage. Gfastats (Formenti et al., 2022) was used to output different assembly statistics of the assemblies.

BlobToolKit and BlobTools2 (Laetsch & Blaxter, 2017), in addition to blobtk were used to visualize assembly statistics. To generate the Hi-C contact map image, the Hi-C reads were mapped to the assemblies using BWA-MEM (Li, 2013) using the same approach as above. Finally, PretextMap was used to create a contact map which was visualized using PretextSnapshot.

To characterize the differences between the two pseudo-haplotypes, we ran a genome alignment using minimap2 (Li, 2017) on the homologous chromosomes. The resulting alignment was processed with the minimap2-included paftools.js producing a report listing the number of insertions, SNPs and indels between the two pseudo-haplotypes.

To demonstrate the duplicated nature of *R. chamaemorus*, we analyzed synteny between hap1 and *Rubus idaeus* (red raspberry cv. ‘Autumn Bliss’) using the R package pipeline GENESPACE (Lovell et al., 2022). *R. idaeus* was chosen as reference due to its close phylogenetic relationship and availability of a high-quality chromosome-scale assembly (Martinussen et al., 2013; Price et al., 2023). To improve orthogroup inference, we included additional *Rubus*: *Rubus occidentalis* (v3.0 - black raspberry) and *Rubus argutus* (‘Hillquist’ v1.0 - blackberry), as well as more distantly related Rosaceae members: *Fragaria vesca* (v4.0.a1 - wild strawberry) and *Rosa chinensis* (‘Old Blush’ v2.0 - Chinese rose). Assemblies, protein sequences and annotations were obtained from the Rosaceae database (https://www.rosaceae.org/). We selected the longest protein isoform as the representative for each gene, along with gene rank orders, to serve as input for GENESPACE. The pipeline was run with the default method, which uses OrthoFinder (Emms & Kelly, 2019) to build orthogroups and MCScanX (Wang et al., 2012) to detect collinear regions. All other parameters were left at default. Synteny blocks between *R. chamaemorus* and *R. idaeus* were then visualized using the plot_riparian function of the GENESPACE package.

Subgenome structure in *R. chamaemorus* was assessed using SubPhaser v1.2.6 (Jia et al., 2022). The analysis was run independently on each haplotype, with default parameters and the 7 sets of 4 homologous chromosomes (tetrads) defined via a configuration file.

The doubletrouble R-package (Almeida-Silva & Van de Peer, 2025) was used to calculate pairwise Ks values between segmental duplicates (i.e. duplicated genes in conserved syntenic blocks) in the hap1 assembly. Since we could not reliably group chromosomes into four subgenomes based on shared k-mer signatures found by SubPhaser (Figure 4A), pairwise Ks values were instead stratified by chromosome tetrads.

To estimate divergence times, Ks values were filtered to exclude values greater than 2 to minimize the effects of substitutional saturation. For each chromosome pair within each tetrad, we calculated the median Ks and derived 95% confidence intervals using non-parametric bootstrapping with 1,000 replicates, implemented using the boot package in R. Comparisons with fewer than 10 valid Ks values were excluded. Divergence time (T) was estimated using the formula: T = Ks / (2 × μ), where μ is the substitution rate, set to a constant 4.13 × 10⁻⁹ substitutions/site/year, a mean herbaceous annual and perennial angiosperm rate reported by Kay et al. (2006). Confidence intervals for divergence time were calculated by applying the same formula to the bootstrap-derived lower and upper bounds of Ks.

To assess potential parental contributors to the *R. chamaemorus* subgenomes, we analyzed mapping bias from available *Rubus* species (plus outgroup *Waldsteinia fragarioides*) using two datasets: Carter et al. (2019) “Target Capture Sequencing Unravels *Rubus* Evolution” (PRJNA510412) and Kates et al. (2024) “Shifts in Evolutionary Lability Underlie Independent Gains and Losses of Root-Nodule Symbiosis in a Single Clade of Plants” (PRJNA1022323, PRJNA1021620, PRJNA1021608). To reduce ambiguity among similar α copies, we retained only the longest α chromosome per tetrad together with the complete β subgenome and aligned reads to this reduced α+β reference with Bowtie2. We kept high-confidence alignments (MAPQ ≥ 30) to limit mismapping across homeologous regions, quantified bias between α and β chromosome pairs as the log2 ratio of read densities (reads per chromosome length) for β versus α [log2(β/α)], and centered each pair so that *R. chamaemorus* had a bias of 0. For each dataset, we projected median species estimates onto the accompanying phylogeny, rooting on *W. fragarioides*: for Carter et al. (2019) the ASTRAL-II exon all-taxa tree (Exon_alltaxa_AstralII.tre), and for Kates et al. (2024) Supplementary Dataset 3 (41467_2024_48036_MOESM6_ESM.txt).

The text in Methods and parts of Results is based on a template we use for all the species we publish in the EBP-Nor project.

## Acknowledgments

This project received data management and infrastructure support from ELIXIR Norway, supported by the Research Council of Norway’s grant 270068, the University of Bergen, the University of Oslo, the Arctic University of Norway in Tromsø, the Norwegian University of Science and Technology and the Norwegian University of Life Sciences: NMBU. The authors acknowledge support from the National Infrastructure for High Performance Computing and resources provided by Sigma2 as well as Data Storage in Norway (project NN8013K) for computational work. The Norwegian Sequencing Centre generated the sequencing data used in this project (http://sequencing.uio.no). We used ChatGPT (temporary chat) for text revision and code refactoring. No AI tool processed study data or created figures; all analyses were performed by the authors.

## Data Availability

All data are available in the European Nucleotide Archive (ENA). The umbrella project for EBP-Nor is BioProject PRJEB98354. Raw PacBio HiFi reads for the drRubCham1 sample (BioSample SAMEA120268085) are deposited under run accessions ERR15658288 and ERR15658311. Illumina Hi-C sequencing reads are available under run accession ERR15658302. The two assemblies are archived separately: pseudo-haplotype 1 under BioProject PRJEB98298 (assembly GCA_976986655) and pseudo-haplotype 2 under BioProject PRJEB98353 (assembly GCA_976986645)

Genome assemblies and gene annotations are also available at **10.5281/zenodo.15773906**

