## Supplementary Material for "The origin of the octoploid cloudberry (*Rubus chamaemorus*) genome is the result of multiple and complex polyploidization events"

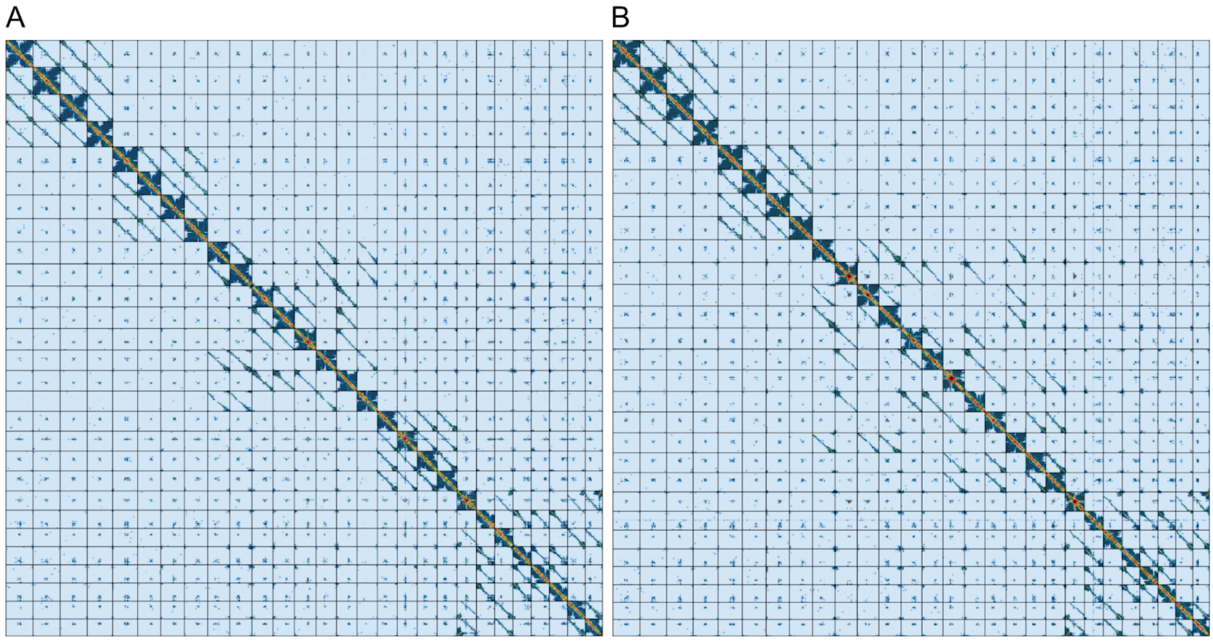


**Supplementary Figure 1: Hi-C contact map of genome assemblies for *R. chamaemorus* hap1 and hap2.** A) Hi-C contact map for hap1. B) Hi-C contact map for hap2. The assemblies are visualized using PreTextSnapshot. Chromosomes are shown in order of size from left to right and top to bottom, therefore scrambling the homologous groups.

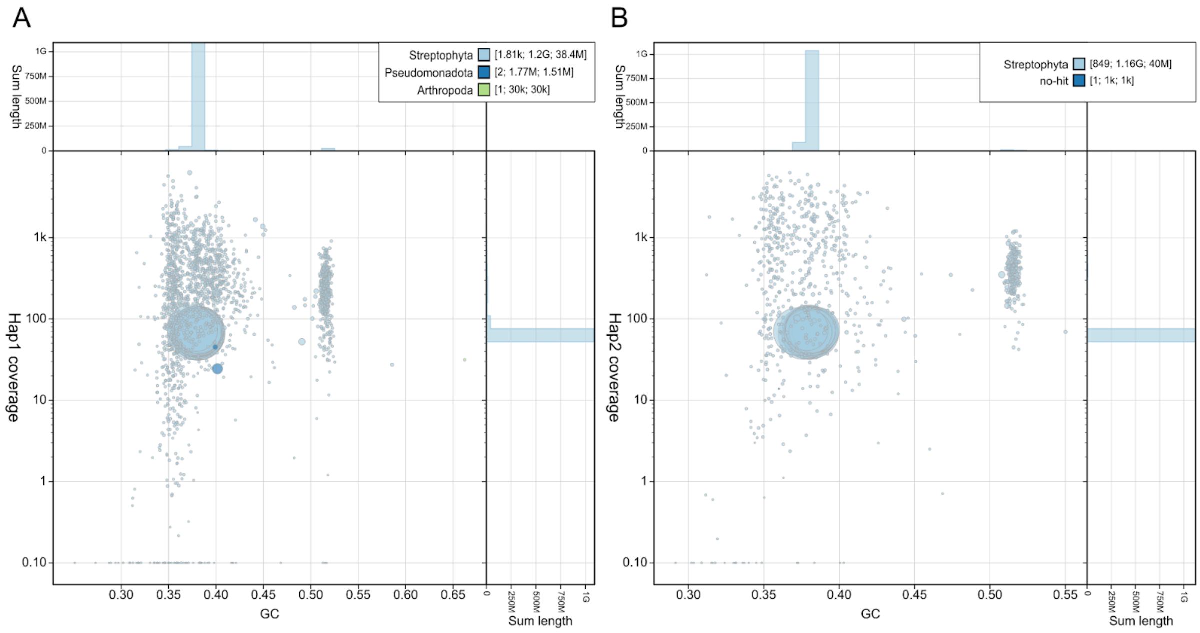


**Supplementary Figure 2: BlobToolKit GC-coverage plots of genome assemblies of *R. chamaemorus* hap1 and hap2.** A) GC-coverage plot for hap1. B) GC-coverage plot for hap2. The scaffolds are coloured by phylum. The size of the circles are in proportion to the length of the scaffolds. Histograms show the distribution of scaffold length sum along each axis.

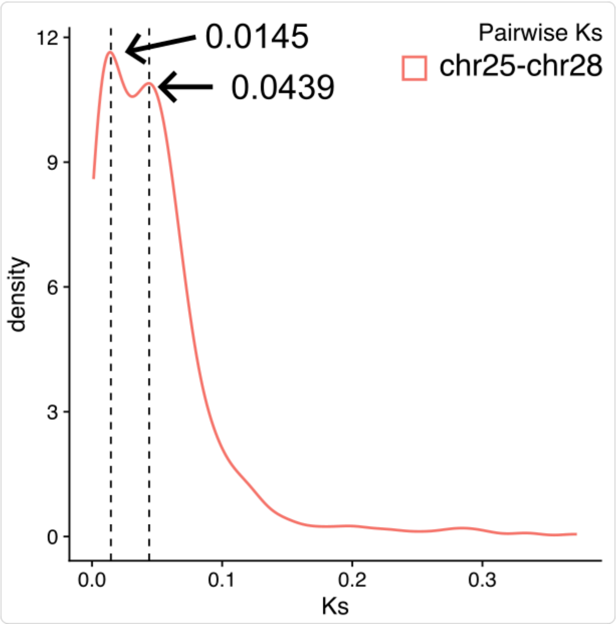


**Supplementary Figure 3: Ks distribution between chromosomes 25 and 28 in Tetrad 7.** Pairwise Ks distribution for chromosome pair 25–28 reveals two distinct peaks, unlike any other combination of chromosome pairs. Vertical dashed lines indicate the estimated Ks peak positions (~0.0145 and ~0.0439).

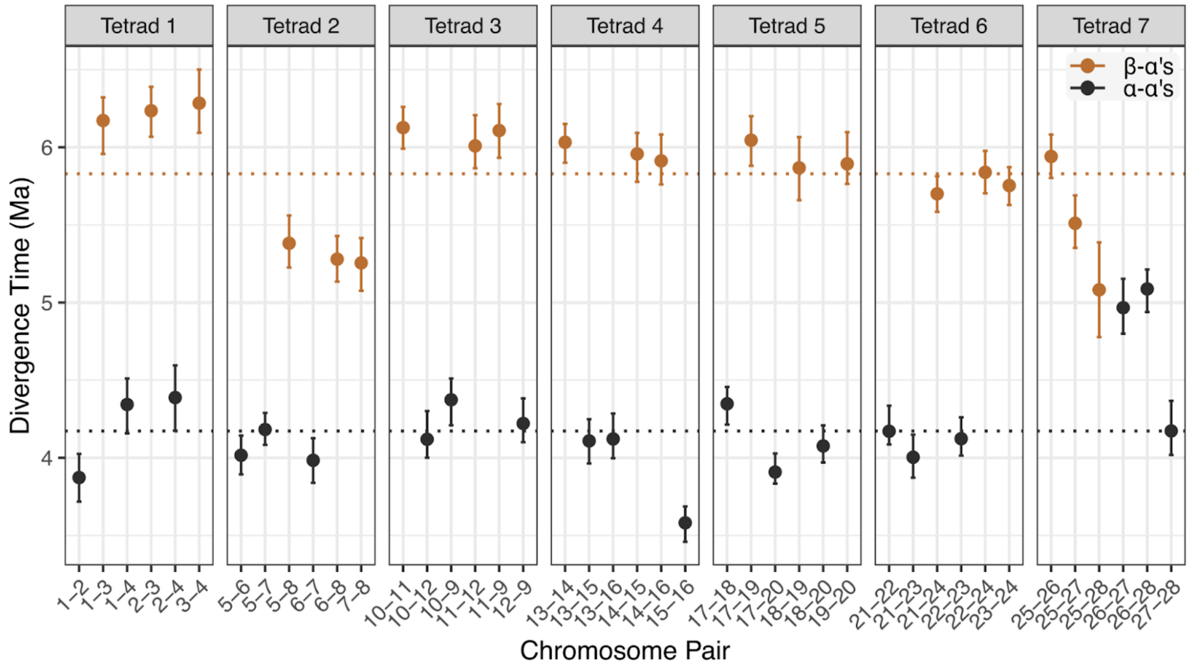


**Supplementary Figure 4. Divergence time estimates by chromosome pair.** Each point represents the estimated divergence time (in million years, calculated with a fixed substitution rate of $4.13 \times 10^{-9}$ substitutions/site/year) for a pair of duplicated chromosomes, based on median Ks values and bootstrap-derived 95% confidence intervals. Chromosome pairs are shown on the x-axis, grouped by tetrad. Orange points and error bars correspond to comparisons between the β-subgenome chromosome and α-subgenome chromosomes (α-β), while dark grey represents comparisons among α-subgenome chromosomes (α-α). Horizontal dotted lines indicate the overall median divergence time for each comparison type (α-β: 5.83 Ma, α-α: 4.17 Ma).

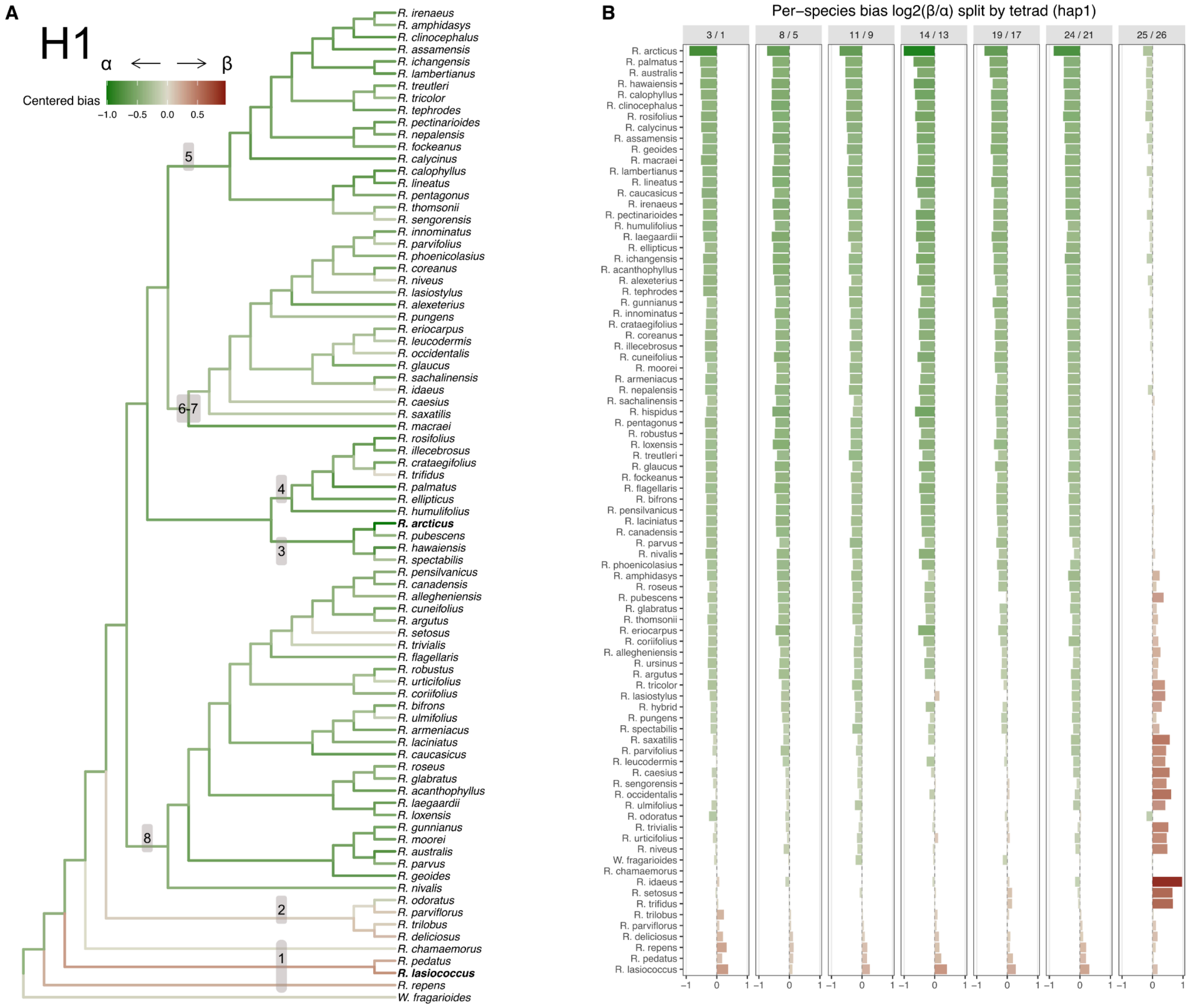


**Supplementary Figure 5: High-confidence read mappings (MAPQ ≥ 30) from Carter et al. (2019) depicting bias of *Rubus* species mapped to hap1 of the reduced α and β subgenome set of *R. chamaemorus*.** Reads from each species are mapped to *R. chamaemorus* haplotype one (hap1). $Log_2$ of the ratio of high-confidence reads mapped to subgenome β over subgenome α, with bias shifted so that *R. chamaemorus* is at 0. A value of 1 indicates a two-fold mapping bias toward subgenome β. Chromosomes from subgenome α: 1, 5, 9, 13, 17, 21, 26; from subgenome β: 3, 8, 11, 14, 19, 24, 25. **A)** Bias projected onto the ASTRAL-II exon all-taxa tree. (Numbers denote groups as defined by Carter et al. (2019, Fig. 2)). The α-biased and β-biased species are marked in bold. **B)** Bias stratified by chromosome pair (tetrad). For each tetrad, taxa are ordered by median bias from most α-biased (most negative) to most β-biased (most positive).

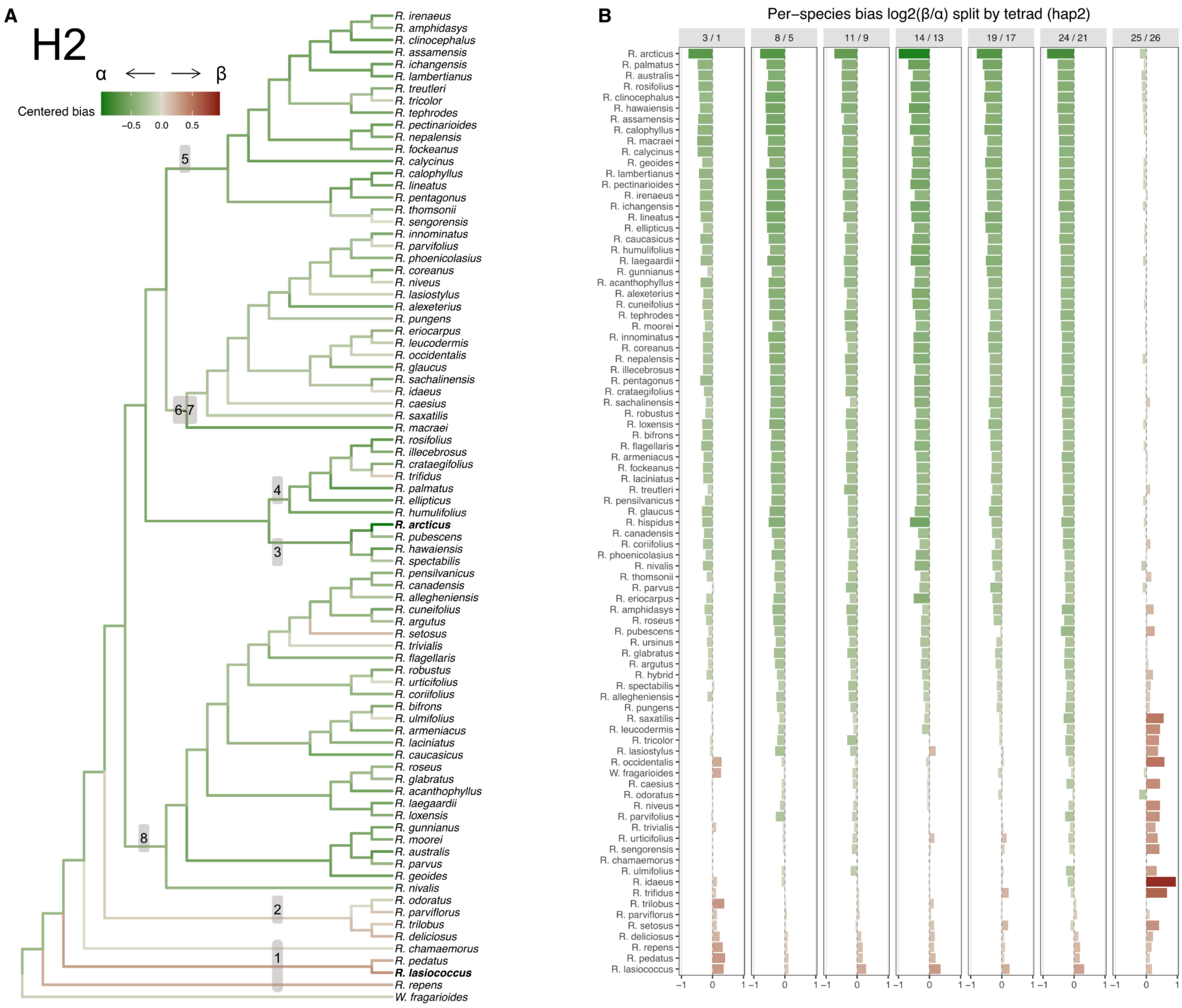


**Supplementary Figure 6: High-confidence read mappings (MAPQ ≥ 30) from Carter et al. (2019) depicting bias of *Rubus species* mapped to hap2 of the reduced α and β subgenome set of *R. chamaemorus*.** Reads from each species are mapped to *R. chamaemorus* haplotype two (hap2). $\text{Log}_2$ of the ratio of high-confidence reads mapped to subgenome β over subgenome α, with bias shifted so that *R. chamaemorus* is at 0. A value of 1 indicates a two-fold mapping bias toward subgenome β. Chromosomes from subgenome α: 1, 5, 9, 13, 17, 21, 26; from subgenome β: 3, 8, 11, 14, 19, 24, 25. **A)** Bias projected onto the ASTRAL-II exon all-taxa tree. (Numbers denote groups as defined by Carter et al. (2019, Fig. 2)). The α-biased and β-biased species are marked in bold. **B)** Bias stratified by chromosome pair (tetrad). For each tetrad, taxa are ordered by median bias from most α-biased (most negative) to most β-biased (most positive).

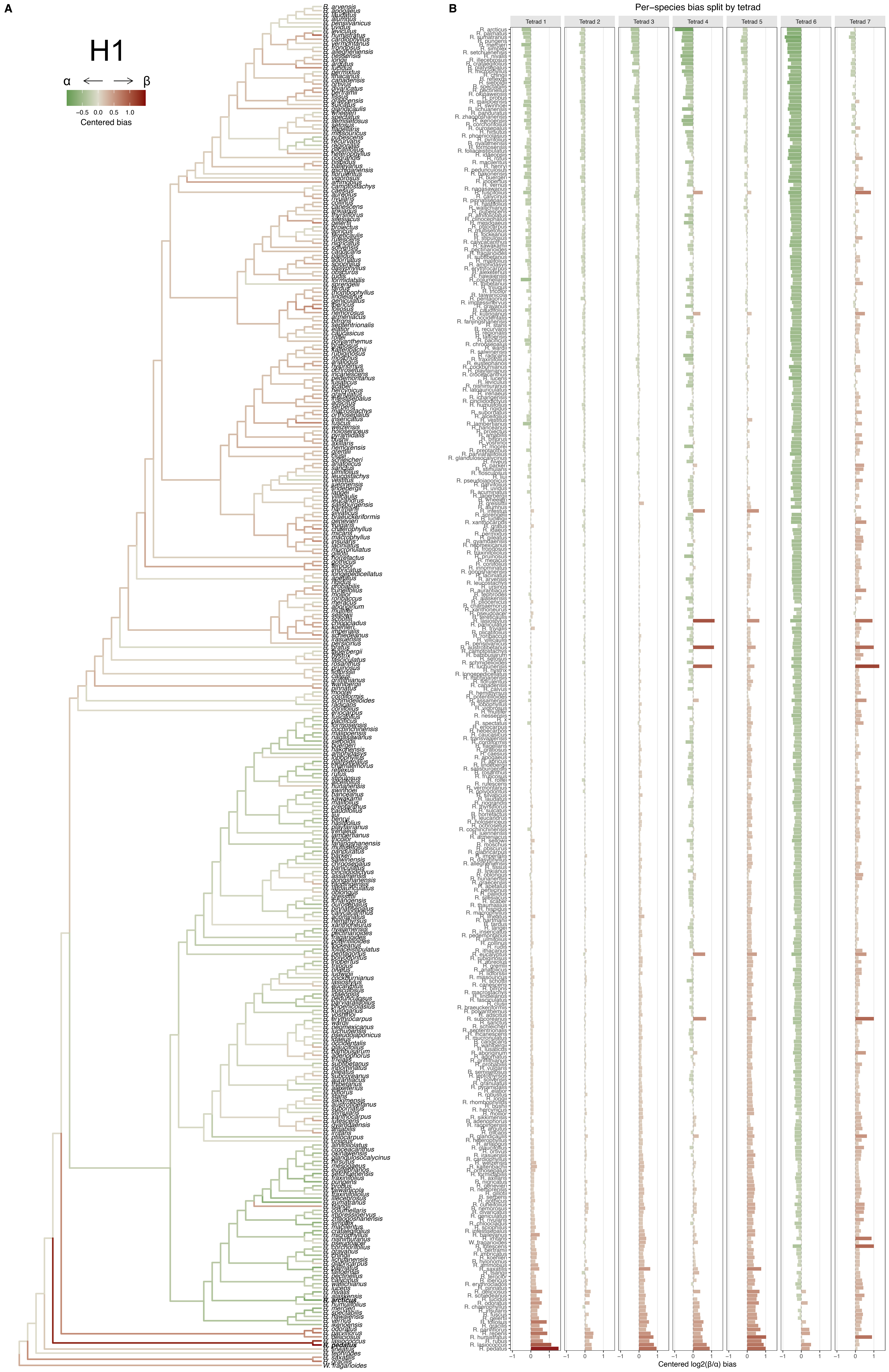
A
H1
α
β
−0.5 0.0 0.5 1.0
Centered bias
B
Per−species bias split by tetrad
Tetrad 1
Tetrad 2
Tetrad 3
Tetrad 4
Tetrad 5
Tetrad 6
Tetrad 7
Centered log2(β/α) bias

**Supplementary Figure 7: High-confidence read mappings (MAPQ ≥ 30) from Kates et al. (2024) depicting bias of *Rubus* species mapped to hap1 of the reduced α and β subgenome set of *R. chamaemorus*.** Reads from each species are mapped to *R. chamaemorus* haplotype one (hap1). $Log_2$ of the ratio of high-confidence reads mapped to subgenome β over subgenome α, with bias shifted so that *R. chamaemorus* is at 0. A value of 1 indicates a two-fold mapping bias toward subgenome β. Chromosomes from subgenome α: 1, 5, 9, 13, 17, 21, 26; from subgenome β: 3, 8, 11, 14, 19, 24, 25. **A)** Bias projected onto the Kates et al. 2024 supplementary dataset 3 phylogeny (41467_2024_48036_MOESM6_ESM.txt). The α-biased and β-biased species are marked in bold. **B)** Bias stratified by chromosome pair (tetrad). For each tetrad, taxa are ordered by median bias from most α-biased (most negative) to most β-biased (most positive).

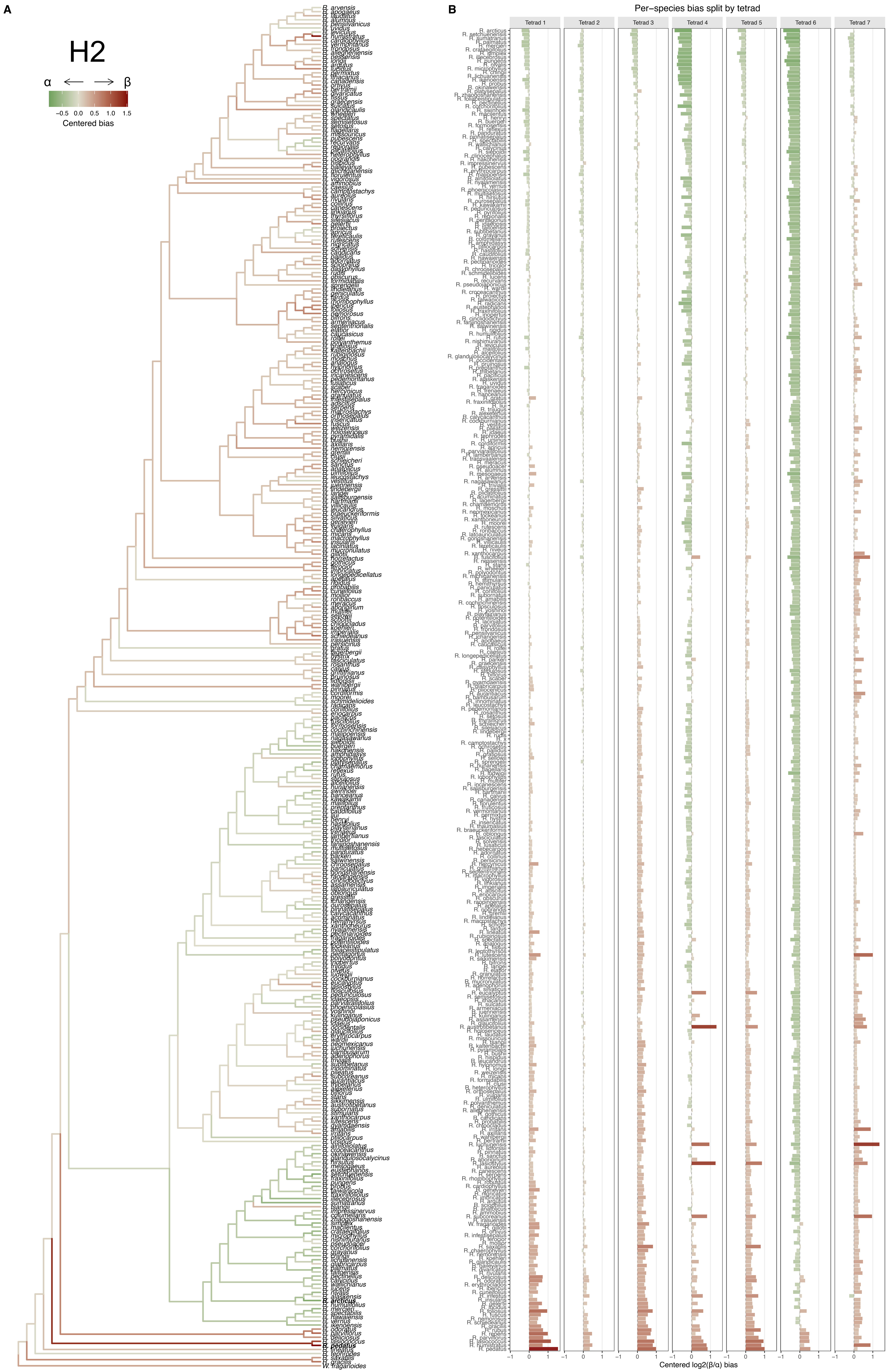
A
H2
α
β
−0.5 0.0 0.5 1.0 1.5
Centered bias
B
Per−species bias split by tetrad
Tetrad 1
Tetrad 2
Tetrad 3
Tetrad 4
Tetrad 5
Tetrad 6
Tetrad 7
Centered log2(β/α) bias

**Supplementary Figure 8: High-confidence read mappings (MAPQ ≥ 30) from Kates et al. (2024) depicting bias of *Rubus* species mapped to hap2 of the reduced α and β subgenome set of *R. chamaemorus*.** Reads from each species are mapped to *R. chamaemorus* haplotype two (hap2). $Log_2$ of the ratio of high-confidence reads mapped to subgenome β over subgenome α, with bias shifted so that *R. chamaemorus* is at 0. A value of 1 indicates a two-fold mapping bias toward subgenome β. Chromosomes from subgenome α: 1, 5, 9, 13, 17, 21, 26; from subgenome β: 3, 8, 11, 14, 19, 24, 25. **A)** Bias projected onto the Kates et al. 2024 supplementary dataset 3 phylogeny (41467_2024_48036_MOESM6_ESM.txt). The α-biased and β-biased species are marked in bold. **B)** Bias stratified by chromosome pair (tetrad). For each tetrad, taxa are ordered by median bias from most α-biased (most negative) to most β-biased (most positive).

**Supplementary table 1**. Software tools: versions and sources

| Software tool | Version | Source |
|---|---|---|
| BlobToolKit | 4.3.2 | https://github.com/blobtoolkit/blobtoolkit |
| blobtk | 0.5.8 | https://github.com/blobtoolkit/blobtk |
| BUSCO | 5.4.7 | https://gitlab.com/ezlab/busco |
| Hifiasm | 0.19.8 | https://github.com/chhylp123/hifiasm |
| KMC | 3.2.1 | https://github.com/refresh-bio/KMC |
| GenomeScope | 2.0 | https://github.com/tbenavi1/genomescope2.0 |
| HiFiAdapterFilt | 2.0.1 | https://github.com/sheinasim/HiFiAdapterFilt |
| PretextView | 1.0.1 | https://github.com/sanger-tol/PretextView |
| PretextMap | 0.1.9 | https://github.com/sanger-tol/PretextMap |
| PretextSnapshot | 0.0.4 | https://github.com/sanger-tol/PretextSnapshot |
| meryl | 1.3.0 | https://github.com/marbl/meryl |
| BWA-MEM | 0.7.17 | https://github.com/lh3/bwa |
| samtools | 1.17 | https://github.com/samtools/samtools |
| YaHS | 1.2a.2 | https://github.com/c-zhou/yahs |

| | | |
|---|---|---|
| FCS-GX | 0.4.0 | https://github.com/ncbi/fcs |
| Merqury | 1.3 | https://github.com/marbl/merqury |
| AGAT | 1.4.0 | https://github.com/NBISweden/AGAT |
| Oatk | 1.0 | https://github.com/c-zhou/oatk |
| miniprot | 0.13 | https://github.com/lh3/miniprot |
| GALBA | 1.0.9 | https://github.com/Gaius-Augustus/GALBA |
| RED | 2018.09.10 | https://github.com/BioinformaticsToolsmith/Red |
| Funannotate | 1.8.17 | https://github.com/nextgenusfs/funannotate |
| EvidenceModeler | 2.1.0 | https://github.com/EVidenceModeler/EVidenceModeler |
| DIAMOND | 2.1.8 | https://github.com/bbuchfink/diamond |
| InterProScan | 5.62-94.0 | https://www.ebi.ac.uk/interpro/search/sequence/ |
| EMBLmyGFF3 | 2.2 | https://github.com/NBISweden/EMBLmyGFF3 |
| Earl Grey | 4.1.1 | https://github.com/TobyBaril/EarlGrey |
| Rapid curation 2.0 | 964d17e997e00c69f25940cf96d3658bda631147 | https://github.com/Nadolina/Rapid-curation-2.0, |
| GENESPACE | 1.3.1 | https://github.com/jtlovell/GENESPACE |

**Supplementary table 2. Alignment comparisons between assemblies of *Rubus chamaemorus* hap1 and hap2. Hap2 aligned to hap1.**

| | |
|---|---|
| Bases in alignment | 731,366,624 |
| Substitutions (%) | 1,775,487 (2.098%) |
| Total # insertions | 149,763 |
| Total # deletions | 149,334 |

| | Insertions | Deletions |
|---|---|---|
| 1bp | 53,769 | 53,297 |
| 2bp | 22,102 | 22,124 |
| [3,50) | 61,606 | 61,547 |
| [50,1000) | 7,110 | 7,023 |
| >=1000 | 5,176 | 5,343 |

**Supplementary table 3: Median Ks distances between tetrads of syntenic chromosomes based on segmentally duplicated genes.**

| Tetrad 1 | chr 1 | chr 2 | chr 3 | chr 4 |
|---|---|---|---|---|
| chr 1 | NA | 0.032 | 0.051 | 0.036 |
| chr 2 | 0.032 | NA | 0.052 | 0.036 |
| chr 3 | 0.051 | 0.052 | NA | 0.052 |
| chr 4 | 0.036 | 0.036 | 0.052 | NA |

| Tetrad 2 | chr 5 | chr 6 | chr 7 | chr 8 |
|---|---|---|---|---|
| chr 5 | 2.239 | 0.033 | 0.035 | 0.045 |
| chr 6 | 0.033 | 1.988 | 0.033 | 0.044 |
| chr 7 | 0.035 | 0.033 | 2.496 | 0.044 |
| chr 8 | 0.045 | 0.044 | 0.044 | 2.069 |

| Tetrad 3 | chr 9 | chr 10 | chr 11 | chr 12 |
|---|---|---|---|---|
| chr 9 | 3.254 | 0.036 | 0.051 | 0.035 |
| chr 10 | 0.036 | 2.181 | 0.051 | 0.035 |
| chr 11 | 0.051 | 0.051 | 2.512 | 0.05 |
| chr 12 | 0.035 | 0.035 | 0.05 | 1.903 |

| | chr 13 | chr 14 | chr 15 | chr 16 |
|---|---|---|---|---|
| chr 13 | NA | 0.05 | 0.034 | 0.034 |

| | | | | |
|---|---|---|---|---|
| chr 14 | 0.05 | NA | 0.049 | 0.049 |
| chr 15 | 0.034 | 0.049 | NA | 0.03 |
| chr 16 | 0.034 | 0.049 | 0.03 | NA |

| | chr 17 | chr 18 | chr 19 | chr 20 |
|---|---|---|---|---|
| chr 17 | NA | 0.036 | 0.05 | 0.032 |
| chr 18 | 0.036 | NA | 0.048 | 0.034 |
| chr 19 | 0.05 | 0.048 | NA | 0.049 |
| chr 20 | 0.032 | 0.034 | 0.049 | NA |

| | chr 21 | chr 22 | chr 23 | chr 24 |
|---|---|---|---|---|
| chr 21 | NA | 0.035 | 0.033 | 0.047 |
| chr 22 | 0.035 | NA | 0.034 | 0.048 |
| chr 23 | 0.033 | 0.034 | NA | 0.048 |
| chr 24 | 0.047 | 0.048 | 0.048 | NA |

| | chr 25 | chr 26 | chr 27 | chr 28 |
|---|---|---|---|---|
| chr 25 | 1.875 | 0.049 | 0.046 | 0.042 |
| chr 26 | 0.049 | 2.006 | 0.041 | 0.042 |
| chr 27 | 0.046 | 0.041 | 2.297 | 0.035 |
| chr 28 | 0.042 | 0.042 | 0.035 | 1.421 |
